# Effective Tubulin Degradation by Rationally Designed Proteolysis Targeting Chimeras

**DOI:** 10.1101/2025.05.22.655572

**Authors:** Alice Maiocchi, Anne-Catherine Abel, Milo Basellini, Helena Pérez-Peña, Zlata Boiarska, Erica E. Ferrandi, Zuzanna Kozicka, Valerio Fasano, Stefano Pieraccini, Graziella Cappelletti, Michel O. Steinmetz, Daniele Passarella, Andrea E. Prota

## Abstract

Proteolysis targeting chimeras (PROTACs) are heterobifunctional molecules that induce the degradation of proteins of interest (POIs) via the ubiquitin-proteasome pathway by recruiting E3 ligases to form a ternary complex with the POI. In this study, we rationally designed and synthesized PROTACs targeting the αβ-tubulin heterodimer, the building block of microtubules (MTs) that are essential for numerous cellular functions and represent important therapeutic targets in cancer and neurodegenerative diseases. Maytansinol, a known MT-destabilising agent, was selected as the POI ligand, functionalised and conjugated to linkers bearing cereblon or Von Hippel-Lindau ligands as E3 ligase recruiters. Four compounds were synthesized and characterized through structural, biophysical and cell biology studies to evaluate their ability to form degradation-prone tubulin-PROTAC-E3 ligase ternary complexes. We confirmed that the PROTACs effectively bind tubulin and recruit E3 ligases. Remarkably, two PROTACs exhibited cellular degradation activity, representing an important advancement in chemically inducing tubulin-E3-ligase interactions. This work integrates rational design, biophysical and structural validation, and cell-based studies to establish a robust framework for developing tubulin-targeting PROTACs, offering significant implications for basic research and therapeutic developments.

## Introduction

Targeted Protein Degradation (TPD) is a revolutionary concept that exploits cellular protein regulation systems to trigger protein degradation^1^, as opposed to conventional target inhibition. In this context, PROteolysis TArgeting Chimeras (PROTACs) are bivalent compounds designed to recruit an E3 ligase, leading to the ubiquitination of the target protein and marking it for degradation^2^. PROTACs feature two ligands connected via a linker moiety that enables efficient binding to both a protein of interest (POI) and E3 ligase targets, ensuring the formation of a ternary POI-PROTAC-E3 ligase complex^3^. This complex ultimately leads to the polyubiquitination of the POI, its release from the ternary complex, and its subsequent degradation by the proteasome. Notably, this mechanism enables sub-stoichiometric activity, with each PROTAC molecule facilitating multiple cycles of the POI ubiquitination^4^. Several E3 ligases have been harnessed in PROTAC technology, with cereblon (CRBN)^5^ and Von Hippel-Lindau (VHL)^6^ being the two most widely utilized due to their accessibility and the large number of identified ligands.

Given our interest in studying MT dynamics modulated by small molecule binders and probes^7, 8, 9^, we here aimed to investigate the PROTAC-mediated degradation of the αβ-tubulin heterodimer, a highly conserved and essential protein in all eukaryotes. Tubulin dimers assemble into dynamic microtubules (MTs) and perform multiple fundamental cellular functions such as structural support, intracellular transport, and force generation in cell division^10^, with a key role in cancer development and neurodegeneration^11, 12^. Since MTs are implicated in such important cellular mechanisms, MT targeting agents (MTAs) have been developed and used very successfully as anticancer drugs^13^, exerting their effects by inhibiting mitosis in rapidly dividing cells, thereby promoting cell senescence and apoptotic cell death. Moreover, MTAs are also increasingly recognized to stimulate immunogenic responses via specific immune cell signaling pathways leading to the activation of dendritic cells through distinct pathways^14, 15^.

Tubulin degradation driven by small molecules is currently under exploration; so far, only a few small molecules have been shown to act as tubulin degraders through the ubiquitin proteasome system (UPS) ^16, 17, 18^, in addition to a series of colchicine-site binders covalently modifying βCys241 with an unspecified mechanism of action^19^. To date, tubulin-targeting PROTACs reported by Gasic et al.^20^ have been shown to lack any degradation activity, while more recent work by Yang et al.^21^, targeting the colchicine binding site, demonstrated proteasomal degradation of tubulin.

In this regard, we here successfully designed and obtained PROTAC-type molecules targeting the tubulin maytansine site, aiming to promote the formation of ternary tubulin-PROTAC-E3 ligase complexes and demonstrate the effective degradation of tubulin driven by PROTACs through the UPS. We aimed to provide structural insight into the ability of a bivalent compound to conjugate an E3 ligase to tubulin, thereby catalysing its degradation.

## Results and Discussion

### Design of bifunctional tubulin degrader molecules

The maytansine site is a surface exposed, shallow pocket of β-tubulin and ligands targeting this site act on MT polymerization by preventing longitudinal tubulin-tubulin interactions, thereby capping the growing MT ends and promoting their disassembly^22^. We recently demonstrated that the binding mode and affinities of C3-substituted maytansinoids are very similar to the one of maytansine and that bulky moieties at C3-position are well tolerated^7, 23^. For this study, we selected maytansinol as the POI ligand to be conjugated at its C3 position with linkers bearing degradation targeting moieties, as its exposed region offers design flexibility and ample space to recruit another protein. As E3 ligase recruiters, thalidomide and lenalidomide for CRBN and deacetyl-VH032 for VHL were considered. We focused on straightforward and biocompatible connecting functional groups and chemistry to construct the heterobifunctional compounds, based on a stable triazole bridge via ’click chemistry’ (Fig. 1). The copper(I)-alkyne/azide cycloaddition (CuAAC) click reaction is highly efficient and regioselective, and widely applicable in the conjugation of various types of molecules, including complex natural products^24, 25^.

**Fig. 1.**
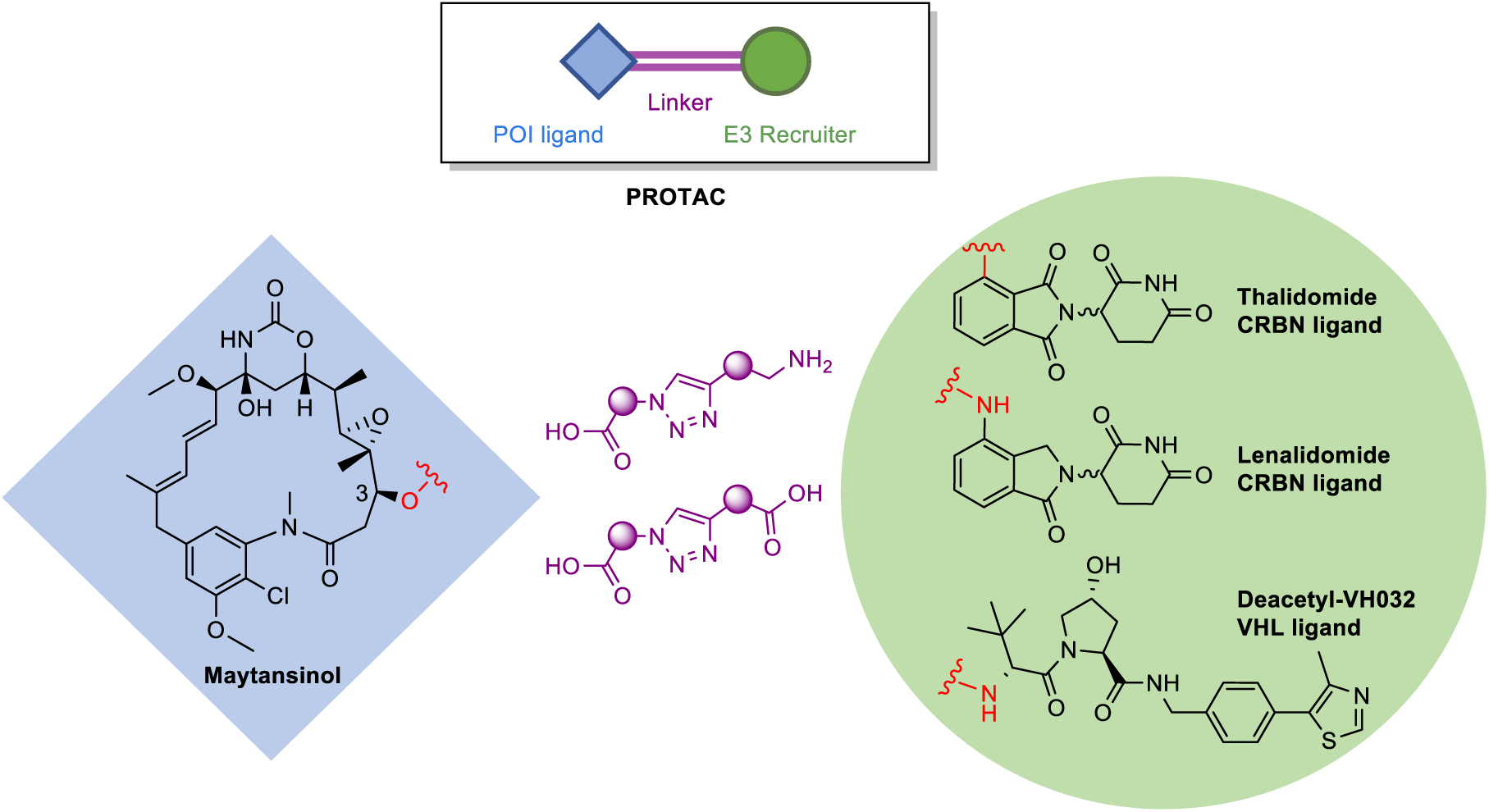
Design of bifunctional tubulin-targeted degraders. Selected moieties for the design of maytansinol-centred PROTACs targeting tubulin and CRBN/VHL E3 ligases.

Due to the fundamental role of both the linker length and chemical nature in PROTAC design and development^26^, we generated a library of 33 possible linkers for evaluation through molecular modelling, varying the triazole position and length of the carbon tails attached to the moieties. In fact, if a linker is too short, it may not enable a dual binding event with the E3 ligase. Instead, an excessively long linker could introduce multiple conformations in the ternary POI-PROTAC-E3 ligase complex, ultimately hindering the ubiquitination event. Only the right linker length and composition would position the two proteins perfectly for POI ubiquitination by the E3 ligase.

Structural models of the β-tubulin-CRBN and β-tubulin-VHL complexes were obtained through protein-protein docking (Fig. 2). A library of potential PROTACs was built, containing maytansinol linked to either lenalidomide, thalidomide or deacetyl-VH032 with one of the 33 proposed linkers. Template docking was employed to ensure that the ligand moieties within each PROTAC aligned with their crystal structure positions. Docking results were first screened based on the computed docking scores. Additionally, a detailed visual inspection was conducted to identify the PROTACs docking poses compatible with proper binding of both maytansinol and the E3 recruiters in their crystallographic binding sites. All the poses where one of the ligands was not properly docked due to the insufficient length of the linker or its tensed conformations were ruled out. At the end of this process, four candidate compounds were identified. Their structures and docked conformations are shown in Fig. 2.

**Fig. 2.**
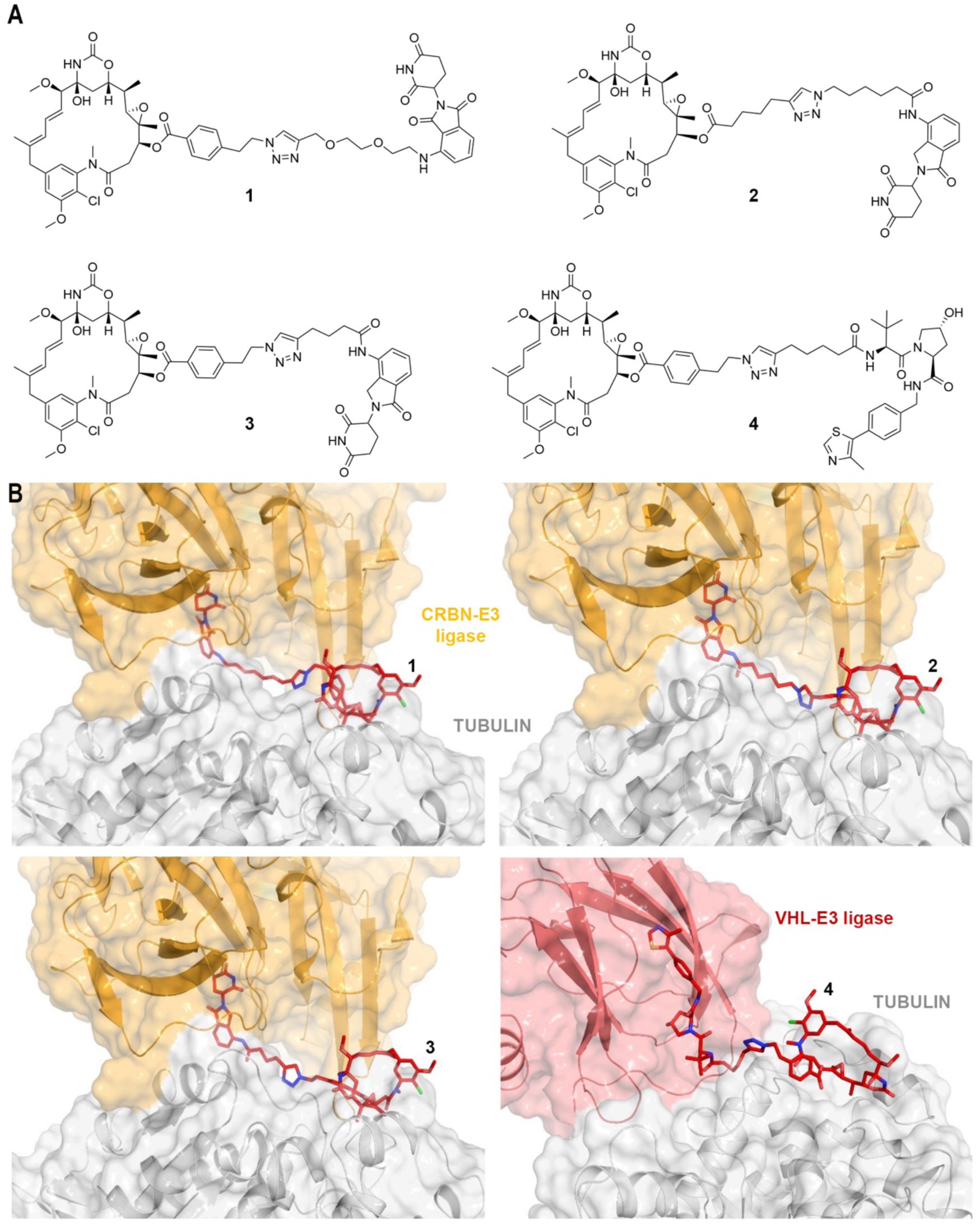
Molecular modelling of bifunctional degrader molecules. (A) 2D chemical structures and (B) docked 3D conformations of four candidate compounds **1**-**4**.

### Synthesis of maytansinol-centered PROTACs

The selected bivalent compounds **1**-**4** can be obtained by a CuAAC reaction between the fragment of an alkyne and an azide providing the corresponding triazole derivatives (Fig. 3). Both the maytansinol-based moieties **5** and **7** can be prepared through esterification starting from maytansinol and 4-(2-bromoethyl)benzoic acid or 6-heptynoic acid. Alkynylamine **6** should be synthesized *via* aryl nucleophilic displacement with fluoro-thalidomide with an alkynyl-PEG_2_-amine; azidoamide **8** should be achieved through an amide condensation between lenalidomide and 6-azidohexanoic acid, while alkynylamides **9** and **10** should be obtained *via* amide condensation between lenalidomide or deacetyl-VH032 and 5- or 6-hexynoic acid.

**Fig. 3.**
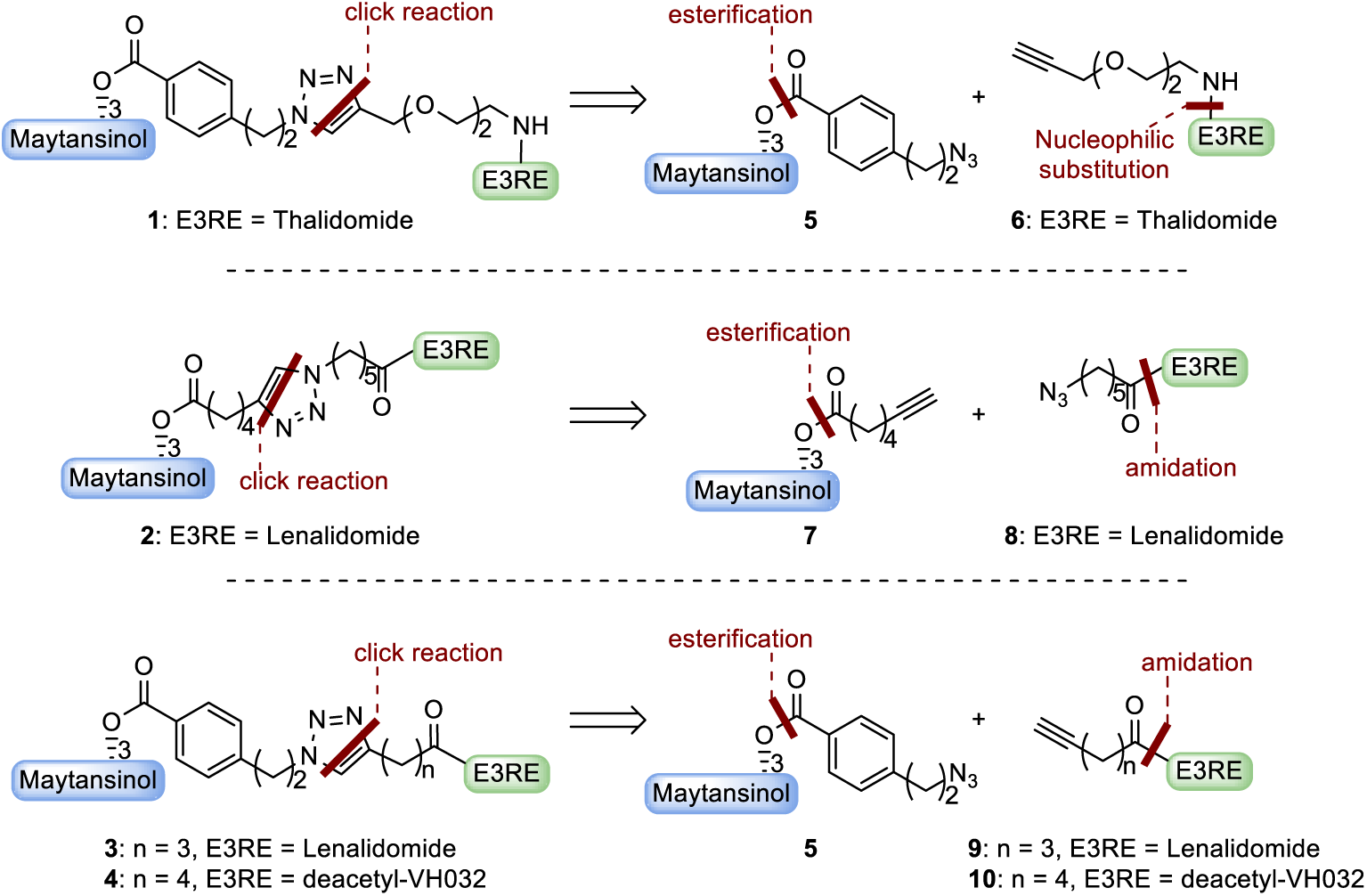
Retrosynthetic pathway for the selected PROTACs **1**-**4**.

Namely, maytansinoids **5** and **7** were obtained by using Steglich esterification (Supplementary Scheme 1) with 6-heptynoic acid or 4-(2-azidoethyl)benzoic acid, which was prepared by treating the corresponding 4-(2-bromoethyl)benzoic acid with sodium azide, with a good overall yield.

Azidoamide **8** and alkynylamide **9** were obtained in quantitative yield by standard amide condensation between lenalidomide and 6-azidohexanoic acid or 5-hexynoic acid, respectively. Similarly, derivative **10** was obtained by amidation between the deacetyl-VH032 and 6-hexynoic acid, employing the same procedure. Conversely, the selected alkynyl-PEG2-amine was reacted with fluoro-thalidomide in basic conditions to carry out an aryl nucleophilic displacement, obtaining intermediate **6** (Supplementary Scheme 2).

Target triazole-connected maytansinol-PROTACs were synthesized under optimized ‘click’ conditions^7^ involving copper sulfate (CuSO_4_) as a catalyst, sodium ascorbate as CuI catalyst regenerator, and DMSO-water as solvent. Target final compounds **1**-**4** were successfully obtained in moderate yields and high purity (Fig. 4), and fully characterized by nuclear magnetic resonance and mass spectrometry analyses.

**Fig. 4.**
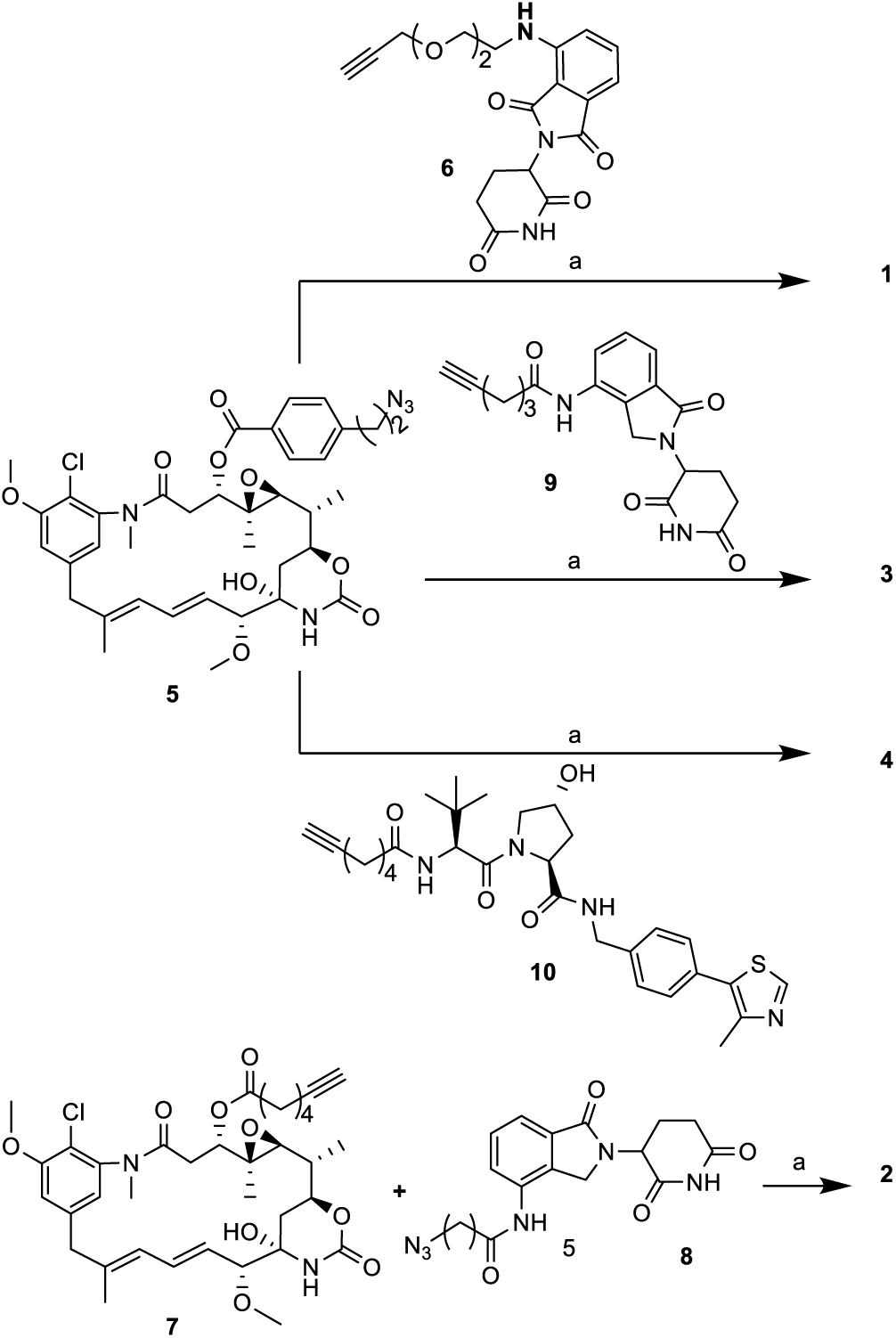
Synthesis of desired PROTAC compounds through ‘click’ chemistry. a) CuSO_4_·5H_2_O, sodium-ascorbate, H_2_O/DMSO 1:2, 3 h, **1**: 37% yield, **2**: 65% yield, **3**: 46% yield, **4**: 41% yield

### Evaluation of the binding of PROTACs to the maytansine site on β-tubulin

To demonstrate the recruitment of the selected POI, our first goal was to confirm that the maytansinol moiety within the PROTAC constructs could bind to the maytansine site of tubulin. As an initial qualitative analysis, we performed thermal shift assays using nanoscale differential scanning fluorimetry (nanoDSF). The analysis required dilution of tubulin into an acidic MES buffer (pH 5.0) to lower the natural high melting point of tubulin from 60 down to 40 °C. Upon addition of PROTACs **1**-**4**, we observed a significant increase in melting temperature of more than 10 °C for all four compounds, indicative of binding to tubulin (Supplementary Fig. S1 and Supplementary Table S1).

Subsequently, we sought to determine the precise binding mode of the compounds by using single crystal X-ray crystallography. Crystals of the T_2_R-TTL complex, composed of two tubulin dimers, the stathmin-like protein RB3, and the tubulin tyrosine ligase (TTL), were grown using the well-established protocols^27, 28^. PROTACs **1-4** were soaked into the crystals at a final concentration of 2.5 or 5 mM for 6 hours. High-resolution X-ray diffraction data (2.2-2.3 Å) were collected, and structures of the complexes were solved (Supplementary Table S4). For all four molecules, ligand-shaped electron density indicated binding to the maytansine site, proving that the E3 recruiter did not prevent binding in the crystal system. In line with our previous observation for long chain linkers^7^, only the central ring of the maytansine scaffold was well defined by the electron density, while the attached linkers and moieties were not visible in any of the obtained tubulin-maytansinoid complex structures. This finding indicates a certain degree of flexibility of the linker and E3-recruiting ligands, likely benefitting the recruitment of ligases. The crystal structures of the compounds were refined to 2.3 Å (**1**, **2**, **3**) and 2.2 Å (**4**) resolution, respectively, and their molecular interactions were analysed (PDB IDs: 9H34, 9H33, 9H32, 9H31). The binding poses of the four tubulin bound molecules are in good agreement with the one of maytansine^22^ (PDB ID: 4TV8). The superpositions of the β-tubulin chains of all the structures revealed no major changes (rmsd**_1_** of 0.18 Å over 385 C_α_ atoms; rmsd**_2_** of 0.17 Å over 372 C_α_ atoms; rmsd**_3_** of 0.19 Å over 383 C_α_ atoms; rmsd**_4_** of 0.18 Å over 367 C_α_ atoms). The macrocycles are bound to the shallow pocket shaped by the side chains of βAsn101, βAsn102, βThr180, βVal181, βVal182, βPhe404, βTrp407, and βTyr408. All hydrogen-bond interactions to the backbone carbonyl of βGly100, the backbone amide of βVal181, and to the side chains of βAsn102 and βLys105 are conserved (Fig. 5). Thus, we concluded that the insertion of the linker moieties at the C3 position of maytansinol is well tolerated in accord with our previous observations^7, 23^.

**Fig. 5.**
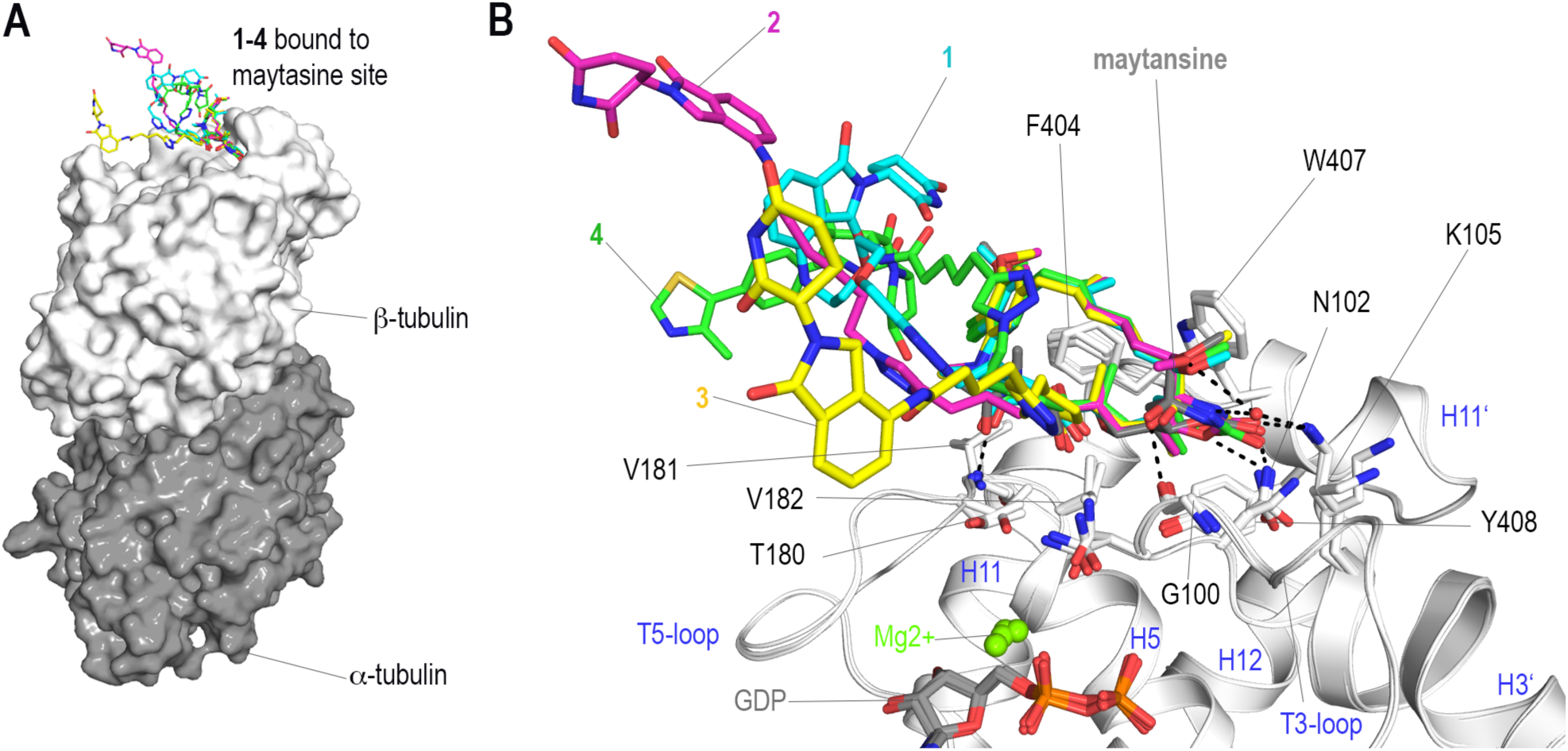
The T2R-TTL complex structures of molecules 1-4. (A) The positions of molecules **1**-**4** bound to the maytansine site on the surface of β-tubulin are shown. The ligands are shown in stick representation and coloured cyan, magenta, yellow, and green, respectively. The α- and β-tubulin monomers are shown in dark grey and white surface representation. (B) Superposition of the complex structures onto the maytansine complex structure (PDB ID 4TV8), with maytansine shown in dark grey sticks. β-Tubulin is shown in ribbon representation and interacting residues are highlighted as sticks. Oxygen, nitrogen, and phosphor atoms are coloured in red, blue, and bright orange, respectively. The guanosine-diphosphate (GDP) molecules are shown in grey sticks and the coordinated magnesium ions as bright green spheres. All residues and secondary structural elements, labelled in black and blue font, belong to β-tubulin and conserved hydrogen bonds are indicated as black dashed lines.

### Evaluation of compounds 1-4 binding to the selected ligases CRBN and VHL

Next, we validated the binding of our PROTAC constructs to the selected E3 ligases again by using nanoDSF experiments. VHL was used in complex with ElonginC (EloC) and ElonginB (EloB; the resuting tripartite complex is called VCB), while CRBN in complex with the damaged DNA–binding protein 1 (DDB1). A clear shift of the melting temperature, indicative of binding to the respective E3 ligase was observed (Supplementary Figs S2 and S3; Supplementary Tables S2 and S3). In contrast, no deviations from the melting point of the reference protein were observed when incubating PROTACs **1-3** with the VCB complex or PROTAC **4** with CRBN (Supplementary Figs S2 and S3; Supplementary Tables S2 and S3).

Next, we sought to determine the crystal structure of the **4**-VCB complex, in order to confirm the binding mode of the ligand and to assess if the maytansinol moiety would be accessible. VCB was incubated with **4** at 1 mM final concentration for 1 hour at room temperature before crystallisation. Crystals were prepared^29^ and X-ray diffraction data were collected (Supplementary Table S4). The crystal structure was solved by molecular replacement and refined to 2.5 Å resolution (PDB ID 9H30). Although a new crystal form with a monoclinic lattice (space group C121) was obtained, differing from the previously reported tetragonal system (PDB ID 4W9H, space group P4_1_22) (Supplementary Fig. S4), the modelled VCB structures were highly similar. The rmsd was 0.30 Å over 121 of 144 C_α_-atoms, with only minor deviations in the EloB subunit and almost perfect agreement over the VHL and EloC chains. Careful analysis of the obtained electron density revealed difference density within the expected binding site allowing the placement of compound **4.** The refined structure superposes well with the known structure of VH032 (PDB ID 4W9H), and the hydrogen bond network is mostly conserved: direct hydrogen bonds are established with VHL residues Tyr98, His110, Ser111, H115 and water-mediated interactions with Tyr112, as well as the interaction with the structural water bound by Val66, Ser68 and Arg69 (Fig. 6).

**Fig. 6.**
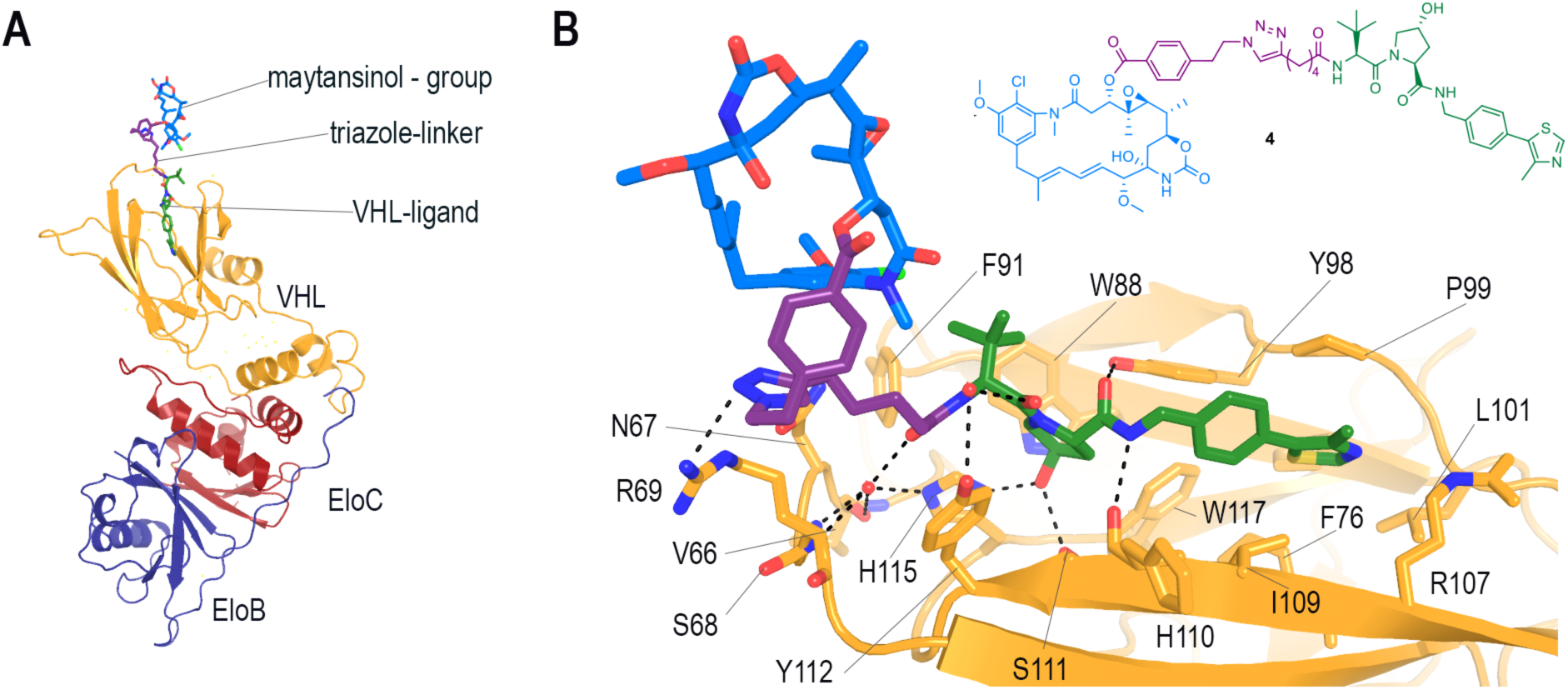
Crystal structure of the VCB-4 complex. (A) Overview of the VCB-**4** complex structure (PDB ID 9H30). The VHL, EloC, and EloB subunits are shown in ribbon representation and are coloured in yellow, red, and dark blue, respectively. Compound **4** is shown in sticks representation. (B) Molecular interactions of **4** with the VHL binding site are shown. The chemical structure of **4** is depicted on the top right and its stick model is colour coded accordingly with the maytansinol, linker, and VHL binding site in blue, dark purple, and green, respectively. Oxygen, nitrogen, and sulphur atoms are coloured in red, blue, and yellow, respectively. Hydrogen bonds are depicted as black dashed lines.

In addition, the triazole linker introduced to couple the maytansinol scaffold to the deacetyl-VH032 establishes one additional hydrogen bond to the side chain of Arg69. Notably, the inversion of the amide bond conformation by connection of the linker and maytansinoid moiety prevents its involvement in a second water mediated hydrogen bond to Tyr112. Surprisingly, the linker and maytansinol moieties were well defined by additional electron density, as the moiety is coordinated at crystal contact sites and thus fixed in a stable conformation (Supplementary Fig. S4), likely imposing the monoclinic crystal form.

### Demonstration of ternary tubulin-PROTAC-ligase complex formation

To investigate whether our PROTACs would facilitate the formation of stable ternary tubulin-PROTAC-E3 ligase complexes, we employed multiangle-light scattering coupled to size exclusion chromatography (SEC-MALS) measurements. Tubulin (10 μM) was mixed with either CRBN (10μM) or VCB complex (10 μM) in the absence or presence of the respective PROTACs **1-3** (10 μM each) or **4** (12 or 24 μM). In the absence of PROTACs only signals corresponding to the reference measurements for tubulin (dubbed peak 3, Fig.7A; dubbed peak 2, Fig.7B), and the isolated E3 ligases CRBN-DDB1τι1 (dubbed peak 2, Fig.7A) and VCB (dubbed peak 3, Fig.7B), were detected with the expected molar masses of 100 kDa, 150 kDa and 40 kDa, respectively (Fig. 7AB). Thus, none of the two ligases show evidence of generally interacting with tubulin dimers without the presence of PROTACs (Supplementary Fig. S5). Upon addition of the PROTACs **1-3** tubulin and CRBN peaks were shifted into a new peak, dubbed peak 1, indicative of formation of a stable tubulin-PROTAC-CRBN complex (Fig. 7A). The extent of complex formation differed between the three compounds tested, with **1** sequestering most of the tubulin into the ternary complex with a calculated molecular weight of 220 kDa. In contrast, PROTACs **2** and **3** (15 μM each) showed a lower potential for promoting the formation of the ternary complex with a significant amount of tubulin remaining as isolated dimer (dubbed peak 3). However, due to both the overlap of the ternary complex with the tubulin peak and the fast exchange kinetics, particularly for compound **3**, the measured molecular weight for the CRBN ternary complexes is not precise. Remarkably, **4** (24 μM) displayed a high potential for promoting ternary complex formation and was able to completely sequester tubulin into the complex with a calculated molecular weight of 125 kDa (dubbed peak1, Fig. 7B). However, the determined size suggests that only the VHL component of the VCB complex was incorporated into the ternary complex, potentially due to the aggregation of EloB and EloC proteins following their dissociation from VHL.

**Fig. 7.**
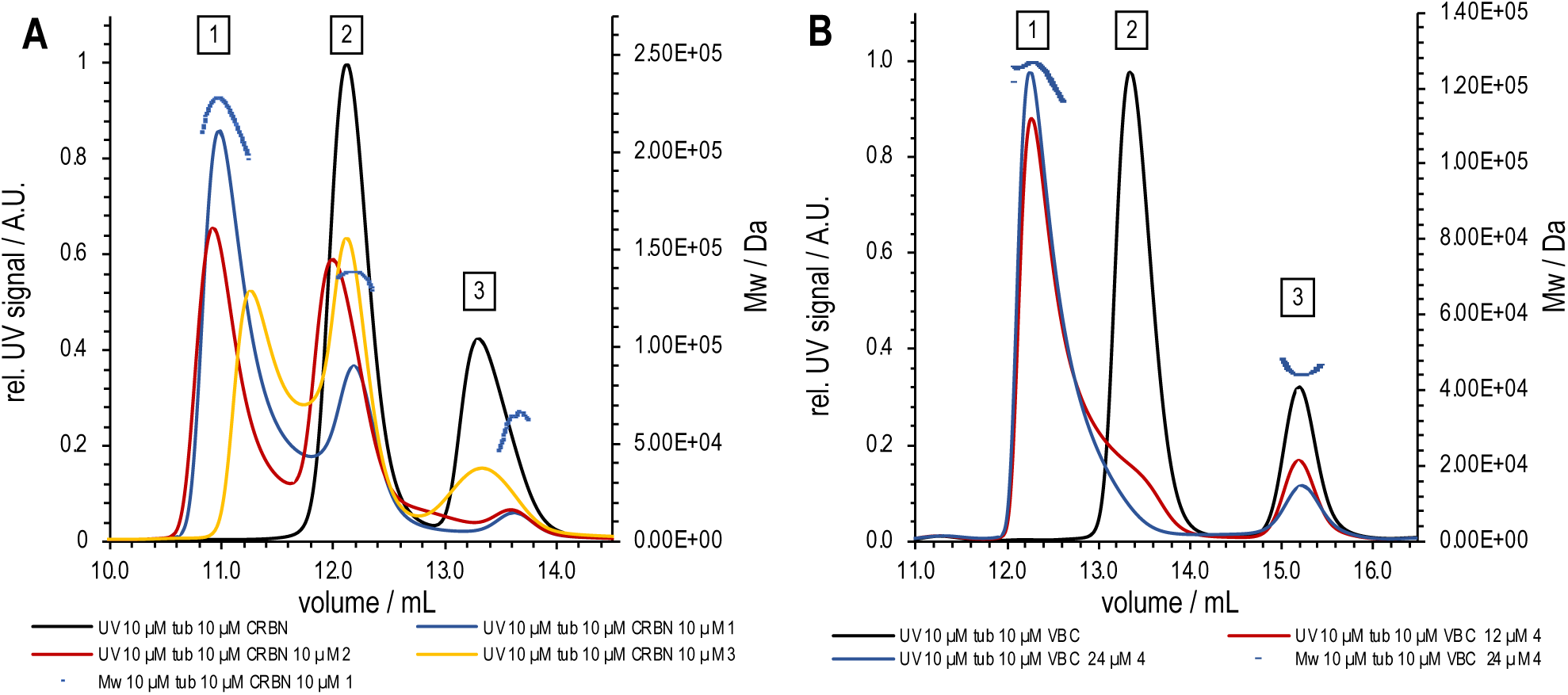
SEC-MALS analysis of ternary tubulin-PROTAC complexes. Chromatograms of tubulin, E3 ligase, and PROTAC mixtures injected onto a SEC column in BRB80 buffer at room temperature are colour coded: UV signals (Y1 axis on the left) are plotted as solid lines, the molecular weights (Y2 axis on the right) as dotted lines. (A) Mixtures of 10 μM tubulin and 10 μM CRBN-DDB1ΔB in the absence (black) and presence of 10 μM of compounds **1** (blue), **2** (red), or **3** (yellow). The molecular weights for molecular species contained in peaks 1, 2, and 3 are shown for the sample containing compound **1**. (B) Mixtures of 10 μM tubulin and 10 μM VHL in the absence (black) and presence of 12 μM (red) or 24 μM (blue) of compound **4**. For the latter concentration, the obtained molecular weights are shown.

### Evaluation of compounds 1-4 cytotoxicity

Firstly, we assessed the cytotoxicity of the compounds using the Hoechst test on CaCo-2 cells and determined IC_50_ values for each compound (Supplementary Fig. S6). As shown in Table 1, compounds **1** and **4** possess an IC_50_ value in the micromolar or submicromolar range, while compounds **2** and **3** appeared to have no toxic effect at the maximum concentration tested (50 μM). This could be due to either low penetrating efficacy, a specific drug resistance mechanism, and/or the intrinsically low toxicity of these two compounds.

**Table 1.** IC_50_ cell viability values determined for compounds 1-4 using the CaCo-2 cell line.

| Compound | IC<sub>50</sub> (μM) | CI 95% (μM) |
| --- | --- | --- |
| **1** | 1.15 x 10<sup>-2</sup> | [0.4 - 4.03] x 10<sup>-2</sup> |
| **2** | > 5 x 10<sup>1</sup> | - |
| **3** | > 5 x 10<sup>1</sup> | - |
| **4** | 2.06 x 10<sup>-1</sup> | [0.7 - 5.58] x 10<sup>-1</sup> |

### Evaluation of in vitro tubulin degradation

The ability of the compounds to degrade tubulin in cells was assessed by western blotting. At maximum confluency, CaCo-2 cells organise into an epithelial-like monolayer and undergo cell cycle arrest through contact inhibition, which helps minimize excessive mitotic activity that could affect the results^30^. All compounds were incubated at increasing concentrations for four hours prior to protein extracts collection, to minimise the contribution of tubulin’s physiological turnover rate^20^. In parallel, the MT destabilising agent batabulin was incubated as a positive control promoting proteasome-dependent tubulin degradation^19^, while maytansinol was also tested to determine if the ligand alone could promote degradation. Compounds **2** and **4** were the most effective in inducing tubulin degradation, showing dose-dependent effects at varying concentration (Fig. 8). In contrast, compounds **1** and **3** did not exhibit any significant impact. Specifically, compound **2** reduced tubulin content by 38% at 10 nM, 56.4% at 100 nM, and 58.9% at 1 μM, but its effect diminished at 10 μM. For compound **4**, tubulin levels were reduced by 33.1% at 100 nM, 46% at 1 μM, and 45% at 10 μM. To verify whether the degradation induced by the compounds was proteasome-mediated, both compounds were tested at their maximum effective concentrations alongside with the proteasome inhibitor MG-132. Co-treatment with 50 μM MG-132 partially prevented tubulin degradation for both compounds, increasing tubulin content by 73.9% for compound **2** and 54.2% for compound **4** relative to the respective single treatments. However, tubulin levels remained 28.5% (compound **2**) and 15.3% (compound **4**) lower than the vector-only control, indicating that the degradation was partially but not entirely proteasome-mediated. Control experiments demonstrated that 10 μM maytansinol failed to induce tubulin degradation, whereas 10 μM batabulin reduced tubulin content by 71.6%. Interestingly, compound **2** remained effective up to a concentration of 1 μM, but its effect diminished when the concentration was increased to 10 μM. This reduced degradation efficiency at higher compound concentration may result from the formation of ineffective binary complexes of the PROTAC, either with the ligase or with tubulin; an inherent entropic drawback also described as ‘hook-effect’^31^ at high PROTAC concentration.

**Fig. 8.**
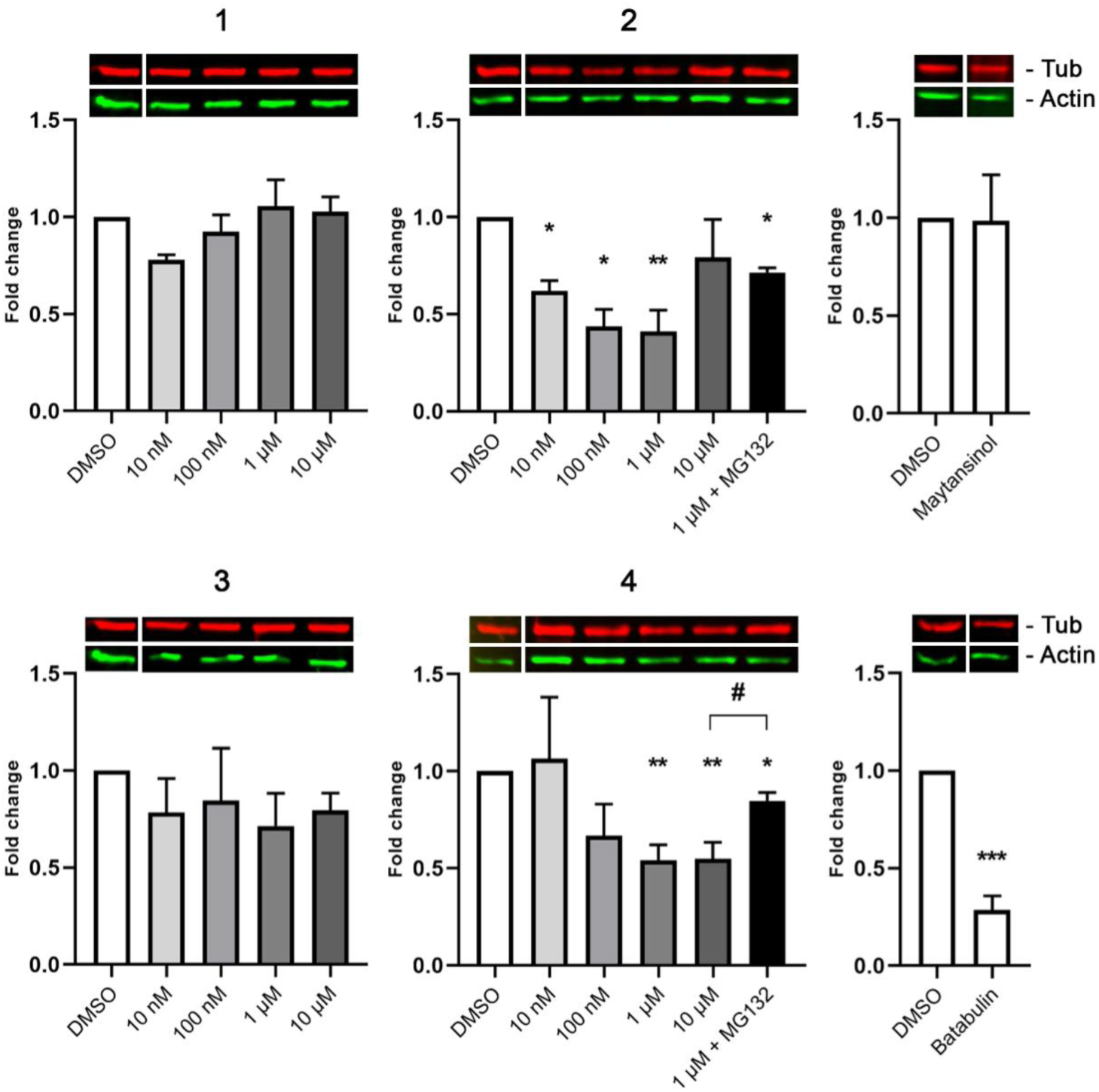
Evaluation of tubulin degradation in CaCo-2 cells. Quantification of relative tubulin levels (red bands); actin (green bands) was used as loading control. Values are expressed as fold change relative to control (0.01% DMSO). Compounds **1** and **3** showed no significant effect. Compound **2** demonstrated a dose-dependent decrease in tubulin levels (10 nM = -38%; 100 nM = -56.4%; 1 μM = -58.9%) plateauing at 1 μM. Co-treatment with 1 μM compound **2** and 50 μM MG-132 increased tubulin levels by 73.9% relative to 1 μM compound **2** alone, with a remaining 28.5% decrease compared to the vector-only control, suggesting partial proteasome-mediated tubulin degradation. Compound **4** also exhibited dose-dependent tubulin degradation (100 nM = -33.1%; 1 μM = -46%; 10 μM = -45%). Co-treatment with 1 μM compound **4** and 50 μM MG-132 restored tubulin levels by 54.2% relative to 10 μM compound **4** alone (# p = 0.016 according to unpaired t-test) but remained 15.3% lower than DMSO control. Control experiments showed that 10 μM maytansinol alone did not induce tubulin degradation, whereas 10 μM batabulin reduced tubulin levels by 71.6%. *p < 0.05; **p < 0.01; ***p < 0.001 versus DMSO, according to One sample t-test.

## Summary

In this work, we rationally designed and synthesised four PROTACs targeting tubulin, leveraging maytansinol as a tubulin binder and CRBN/VHL as E3 ligases. Biophysical and structural characterization confirmed the ability of all compounds tested to bind and recruit tubulin. Additionally, binding to VHL and CRBN was validated. Crucially, we provided evidence for the formation of the ternary tubulin–PROTAC–E3 ligase complex, with two compounds (**2** and **4**) showing tubulin degradation via the proteasome in cell-based assays. Putative PROTAC-induced ubiquitination models mediated by VHL and CRBN E3 ligases identified four exposed lysine residues on the β-tubulin surface (Fig.9) as candidate ubiquitination sites. Two of these lysine residues are highly conserved across tubulin isotypes (βK176 and βK218; residue numbering according to the α-tubulin-based definitions given in^32^), while the other two vary in TUBB1B (βK299R, βK389N), TUBB6 (βK299R), and TUBB8B (βK389T). These models thus underscore the potential of tailored tubulin-targeting PROTACs for studying isotype-specific tubulin degradation.

**Fig.9.**
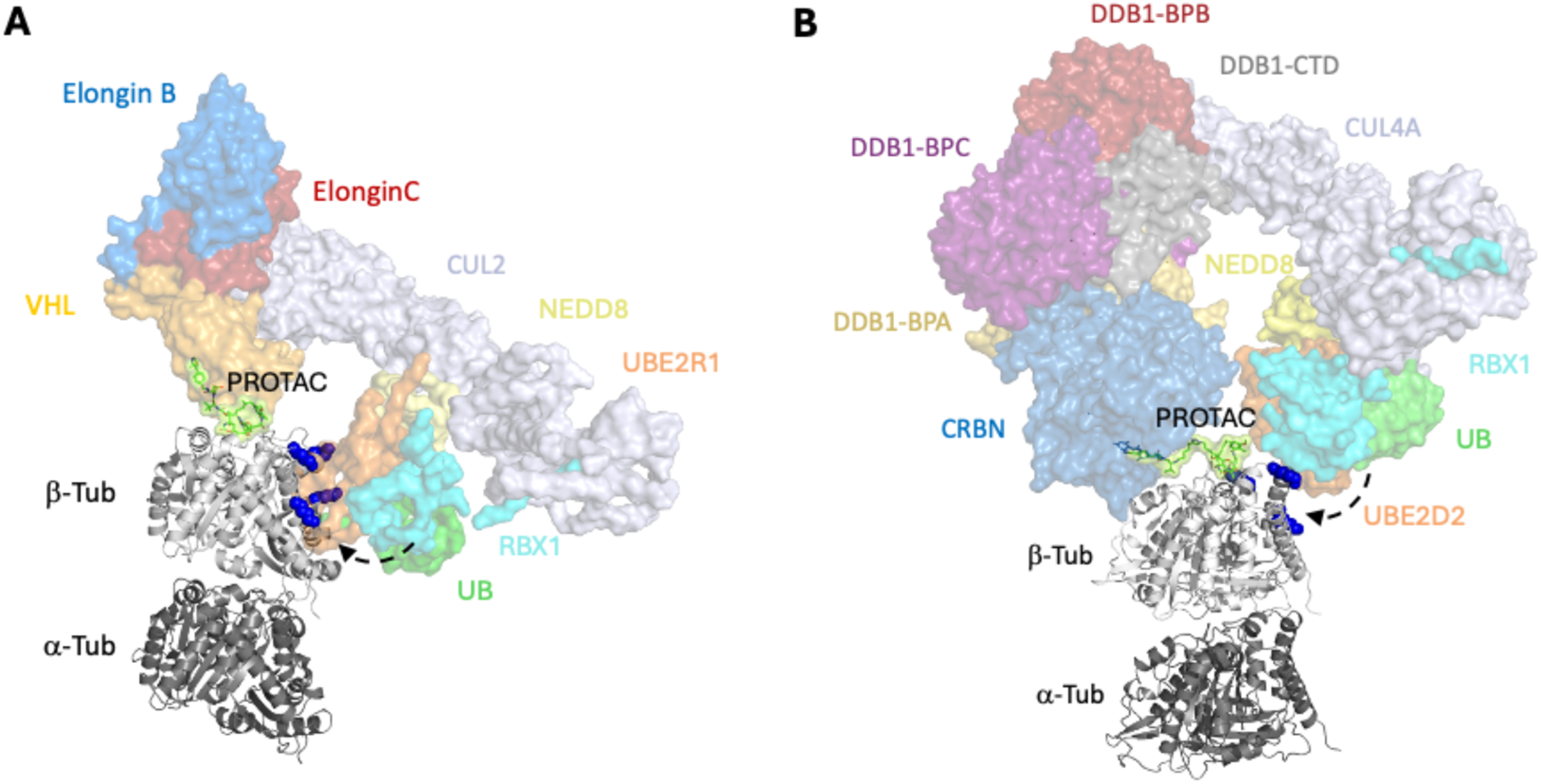
Proposed models of PROTAC-driven tubulin ubiquitination. (**A**) VCB-cullin 2 pathway: Model of a putative functional tubulin-PROTAC-VCB-Cul2-RBX1-UBE2R1-NEDD8-Ub complex. The model was constructed by superimposing the maytansine core heterocycles from the crystal structures of VCB-PROTAC **4** (PDB ID 9H30) and tubulin-PROTAC **4** (PDB ID 9H31) described in this study. The VHL-cullin 2 RING E3 ligase complex (PDB ID 8RX0) was then aligned to the VCB subunit. (**B**) CRBN-cullin 4A pathway: Model of a putative functional tubulin-PROTAC-CRBN-DDB1-Cul4A-RBX1-UBE2D2-NEDD8-Ub complex. This assembly was built starting with the alignment of the lenalidomide isoindole moiety from the cryo-EM structure of CRBN-DDB1 (PDB ID 4CI2) to the corresponding feature in the tubulin-PROTAC **2** complex (PDB ID 9H33). Subsequent superimpositions incorporated the remaining components, including Cul4A-RBX1 (PDB ID 4A0K) and RBX1-UBE2D2-NEDD8-Ub (PDB ID 6TTU). In both proposed models, the ubiquitination machinery aligns with the same interface on β-tubulin, exposing four lysine residues (βLys176, βLys218, βLys299, and βLys389, highlighted as dark blue spheres) as candidate ubiquitination sites. The individually modelled protein subunits are displayed as coloured, semi-transparent surfaces and labelled for clarity.

This study establishes a robust PROTAC development process, combining rational design, synthesis, and *in vitro* and *in cell* validation, to achieve degradation of a challenging target like tubulin. Our findings highlight PROTACs as a powerful tool for modulating tubulin homeostasis, offering a novel approach to next-generation MTAs. These results pave the way for innovative strategies to overcome chemotherapy resistance by efficiently degrading tubulin in cancer cells.

## Methods

### Molecular modelling

All computational protein:protein and protein:ligand docking simulations were conducted using the ICM-Pro software suite (Molsoft LLC, San Diego, CA). Protein structures were retrieved from the Protein Data Bank (PDB). The selected 3D structures were: PDB ID 4TV8^22^, chain D, with the crystal structure of maytansine bound to β-tubulin; PDB ID 5FQD^33^, chain B, containing the crystal structure of lenalidomide bound to CRBN; PDB ID 4W9F^29^, chain I, containing the crystal structure of a VHL recruiter bound to VHL. Protein-protein docking was performed using the Fast Fourier Transform (FFT) method as implemented in ICM-Pro^34^. The following pairs of proteins were docked: β-tubulin-CRBN and β-tubulin-VHL. A library of potential PROTACs was constructed using ICM-Pro, wherein 2D structures were generated containing maytansinol linked to lenalidomide, maytansinol linked to thalidomide, and maytansinol linked to VHL recruiter. These 2D structures were subsequently processed into 3D through the ICM-pro template docking approach. Template docking was employed to ensure that the ligand moieties within each PROTAC aligned with their crystal structure positions. The “fuzzy” option in the ICM-Pro template docking approach was used to identify the maximum common substructure between the template and input ligands.

### Chemical synthesis

Unless otherwise stated, reagents were purchased from suppliers (Sigma-Aldrich, TCI, Fluorochem) and used without further purification. All solvents were of reagent grade or HPLC grade. All reactions were carried out in oven-dried glassware and dry solvents, under nitrogen and were monitored by glasses TLC on silica gel (Merck precoated 60F254 plates), with detection by UV light (254 nm), or by TLC stains. Products were purified by flash column chromatography, using silica gel Merk 60 (230-400 mesh) as stationary phase. ^1^H NMR and ^13^C NMR spectra were recorded on a Bruker Avance Spectrometer 400 MHz using commercially available deuterated solvents at room temperature. Chemical shifts are reported in parts per million (ppm), compared to TMS as an internal standard. Multiplicities in ^1^H-NMR are reported as follow: s – singlet, d – doublet, t – triplet, m – multiplet, br s= broad. Data for ^13^C NMR are reported in terms of chemical shift (δ/ppm).

#### Synthesis of 4-(2-azidoethyl)benzoic acid

NaN_3_ (91 mg, 1.4 mmol, 1.1 eq) was added to a solution of 4-(2-bromoethyl)benzoic acid (300 mg, 1.3 mmol, 1.0 eq) in DMSO (5.2 mL). The mixture was stirred at room temperature overnight, then extracted with EtOAc (4 x 20 mL). The organic phase was washed with brine (1 x 20 mL), dried over anhydrous Na_2_SO_4_ and concentrated under reduced pressure, giving the desired compound (200 mg, 1 mmol, 80% yield) as a white solid, without further purification. ^1^H NMR (400 MHz, CDCl_3_) δ 8.08 (d, *J* = 8.3 Hz, 2H), 7.34 (d, *J* = 8.3 Hz, 2H), 3.56 (t, *J* = 7.1 Hz, 2H), 2.97 (t, *J* = 7.1 Hz, 2H). Spectroscopic data are consistent with that reported in the literature^7^.

#### Synthesis of 6-azidohexanoic acid

NaN_3_ (108 mg, 1.7 mmol, 1.1 eq) was added to a solution of 6-bromohexanoic acid (300 mg, 1.5 mmol, 1 eq) in DMSO (6.1 mL). The mixture was stirred at room temperature for 20 hours. Then, water (10 mL) was added, and the solution was extracted with EtOAc (4 × 10 mL). The organic layer was washed with brine (1 x 10 mL), and the combined organic layers were dried over anhydrous Na_2_SO_4_ and concentrated under reduced pressure, providing the product (205 mg, 1.3 mmol, 85% yield) as white solid. ^1^H NMR (400 MHz, CDCl_3_) δ 3.36 (t, *J* = 6.9 Hz, 2H), 2.32 (t, *J* = 7.4 Hz, 2H), 1.66-1.56 (m, 4H), 1.45-1.38 (m, 2H). Spectroscopic data are consistent with that reported in the literature^7^.

*Synthesis of **5**:* DMAP (65 mg, 0.53 mmol, 3 eq) and 4-(2-azidoethyl)benzoic acid (102 mg, 0.53 mmol, 3.0 eq) were added to a solution of maytansinol (100 mg, 0.18 mmol, 1.0 eq) in dry DCM (0.89 mL) at room temperature under a nitrogen atmosphere. Then, a 1.3 M solution of DCC (121 mg, 0.58 mmol, 3.3 eq) in dry DCM was slowly added and the mixture was stirred at room temperature for 5 hours (TLC moritoring: DCM/MeOH 95:5). Successively, DCU was filtered off using cold DCM and the organic phase was washed with brine (3 x 10 mL), dried with anhydrous Na_2_SO_4_, filtered, and concentrated under reduced pressure. The crude was purified by flash column chromatography (eluent: DCM/MeOH 98:2) to afford **5** as a white solid (60 mg, 0.081 mmol, 45% yield). ^1^H NMR (400 MHz, CDCl_3_) δ 8.02 (d, *J* = 7.5 Hz, 2H), 7.39 (d, *J* = 7.5 Hz, 2H), 7.04 (d, *J* = 1.8 Hz, 1H), 6.85 (d, *J* = 1.8 Hz, 1H), 6.29 (dd, *J* = 15.4, 11.0 Hz, 1H), 6.14 (s, 1H), 5.89 (d, *J* = 11.0 Hz, 1H), 5.02 (dd, *J* = 12.0, 3.3 Hz, 1H), 4.82 (dd, *J* = 15.4, 9.1 Hz, 1H), 4.25 (ddd, *J* = 12.5, 10.5, 2.0 Hz, 1H), 4.00 (s, 3H), 3.59 (td, *J* = 7.0, 2.1 Hz, 2H), 3.48 (d, *J* = 12.0 Hz, 1H), 3.33 (d, *J* = 9.1 Hz, 1H), 3.23 (s, 3H), 3.20 (d, *J* = 12.0 Hz, 1H) 3.18 (s, 3H), 3.12 (d, *J* = 9.6 Hz, 1H), 3.01 (t, *J* = 7.0 Hz, 2H), 2.86 (bs, 1H), 2.75 (dd, *J* = 14.4, 12.0 Hz, 1H), 2.32 (dd, *J* = 14.4, 3.3 Hz, 1H), 1.67 (s, 3H), 1.59 (dt, *J* = 12.5, 2.0 Hz, 1H), 1.50 – 1.43 (m, 1H), 1.29 (d, *J* = 6.4 Hz, 3H), 1.18 - 1.12 (m, 1H), 0.84 (s, 3H). HRMS (ESI^+^) *m/z* [M+Na]^+^ 760.2732; calculated for C_37_H_44_ClN_5_O_9_Na: 760.2725. Spectroscopic data are consistent with that reported in the literature^23^.

*Synthesis of **7**:* DMAP (97 mg, 0.80 mmol, 3 eq) and heptynoic acid (0.100 mL, 0.80 mmol, 3.0 eq) were added to a solution of maytansinol (150 mg, 0.27 mmol, 1.0 eq) in dry DCM (1.30 mL) at room temperature under a nitrogen atmosphere. Then, a 1.3 M solution of DCC (181 mg, 0.88 mmol, 3.3 eq) in dry DCM was slowly added and the mixture was stirred at room temperature for 5 hours (TLC moritoring: DCM/MeOH 95:5). Successively, DCU was filtered off using cold DCM and the organic phase was washed with brine (3 x 10 mL), dried with anhydrous Na_2_SO_4_, filtered, and concentrated under reduced pressure. The crude was purified by flash column chromatography (eluent: DCM/MeOH 98:2) to afford **7** as a white solid (62 mg, 0.092 mmol, 35% yield). ^1^H NMR (400 MHz, CDCl_3_) δ 6.84 (d, *J* = 1.8 Hz, 1H), 6.79 (d, *J* = 1.8 Hz, 1H), 6.44 (dd, *J* = 15.5, 11.0 Hz, 1H), 6.25 (s, 1H), 6.16 (d, *J* = 11.0 Hz, 1H), 5.49 (dd, *J* = 15.5, 8.9 Hz, 1H), 4.89 (dd, *J* = 11.9, 3.0 Hz, 1H), 4.25 (ddd, *J* = 11.3, 10.5, 2.0 Hz, 1H), 3.99 (s, 3H), 3.56 – 3.46 (m, 2H), 3.36 (s, 3H), 3.21 (d, *J* = 13.0 Hz, 1H), 3.18 (s, 3H), 2.89 (d, *J* = 9.7 Hz, 1H), 2.55 – 2.46 (m, 2H), 2.42 – 2.34 (m, 1H), 2.26 (td, *J* = 6.9, 2.6 Hz, 2H), 2.19 (dd, *J* = 13.9, 2.9 Hz, 1H), 1.95 (t, *J* = 2.6 Hz, 1H), 1.81 (p, *J* = 7.6 Hz, 2H), 1.69 (s, 3H), 1.66 – 1.54 (m, 3H), 1.50 – 1.43 (m, 1H), 1.28 (d, *J* = 6.5 Hz, 3H), 1.26 – 1.20 (m, 1H), 0.83 (s, 3H). HRMS (ESI^+^) *m/z* [M+Na]^+^ 695.2710; calculated for C_35_H_45_ClN_2_O_9_Na: 695.2711. Spectroscopic data are consistent with that reported in the literature^23^.

*Synthesis of **6***: 2-(2-(prop-2-yn-1-yloxy)ethoxy)ethan-1-amine (0.085 mL, 0.60 mmol, 1.1 eq) was added to a stirred solution of F-thalidomide (150 mg, 0.54 mmol, 1.0 eq) in dry DMF (2.7 mL) and DIPEA (0.190 mL, 1.1 mmol, 2.0 eq), under nitrogen atmosphere. The reaction mixture was stirred at 90 °C for 5 h and monitored by TLC (eluent: DCM/acetone 9:1). After reaction completion, the mixture was cooled to room temperature, poured into water and extracted with EtOAc (4 x 10 mL). the combined organic layer was washed with brine (1 x 10 mL), dried over anhydrous Na_2_SO_4_ and concentrated under reduced pressure. The crude fluorescent oil was purified by flash column chromatography (eluent: DCM/EtOAc 85:15) to give **6** (106 mg, 0.27 mmol, 49% yield) as a yellow powder. ^1^H NMR (400 MHz, CDCl_3_) δ 8.50 (s, 1H), 7.47 (dd, *J* = 8.5, 7.1 Hz, 1H), 7.08 (d, *J* = 7.1 Hz, 1H), 6.91 (d, *J* = 8.5 Hz, 1H), 4.91 (dd, *J* = 11.8, 5.3 Hz, 1H), 4.19 (d, *J* = 2.4 Hz, 2H), 3.73 – 3.67 (m, 6H), 3.47 (t, *J* = 5.4 Hz, 2H), 2.85 – 2.71 (m, 3H), 2.43 (t, *J* = 2.4 Hz, 1H), 2.14 – 2.06 (m, 1H). ^13^C NMR (101 MHz, CDCl_3_) δ 171.5, 169.4, 168.7, 167.7, 146.9, 136.1, 132.6, 116.9, 111.8, 110.4, 74.8, 70.5, 69.7, 69.2, 58.6, 48.9, 42.5, 31.5, 22.8 (CH of the triple bond not detected). HRMS (ESI^+^) *m/z* [M+H]^+^ 400.1522; calculated for C_20_H_22_N_3_O_6_: 400.1509.

*Synthesis of **8**:* Lenalidomide (100 mg, 0.386 mmol, 1.0 eq) and 6-azidohexanoic acid (91 mg, 0.579 mmol, 1.5 eq) were dissolved in dry DMF (3.2 mL) stirring under nitrogen atmosphere. HATU (147 mg, 0.386 mmol, 1.0 eq) and DIPEA (150 mg, 1.16 mmol, 3.0 eq) were added and the mixture was stirred overnight at room temperature (TLC monitoring: DCM/MeOH 9:1). Then, water and sat. aqueous solution of NH_4_Cl were added, and the solution was extracted with EtOAc (5 x 10 mL). the combined organic layers were dried over Na_2_SO_4_, filtered, and concentrated under reduced pressure. The crude was purified by flash column chromatography (eluent mixture: DCM/MeOH from 95:5 to 9:1) to obtain **8** (74 mg, 0.184 mmol, 48% yield) as a white solid. ^1^H NMR (400 MHz, pyridine-*d_5_*) δ 10.66 (s, 1H), 8.00 (d, *J* = 7.8 Hz, 1H), 7.89 (d, *J* = 7.8 Hz, 1H), 7.46 (t, *J* = 7.8 Hz, 1H), 5.64 (dd, *J* = 13.4, 5.0 Hz, 1H), 4.85 - 4.69 (m, 2H), 3.13 (t, *J* = 6.8 Hz, 2H), 2.96-2.79 (m, 2H), 2.58 (t, *J* = 7.5 Hz, 2H), 2.34 (qd, *J* = 13.0, 4.7 Hz, 1H), 2.10-2.05 (m, 1H), 1.81 (t, *J* = 7.5 Hz, 2H), 1.48-1.43 (m, 2H), 1.42-1.36 (m, 2H). ^13^C NMR (101 MHz, pyridine-*d_5_*) δ 173.5, 172.0, 169.5, 135.3, 134.4, 129.5, 126.3, 120.6, 52.9, 51.7, 48.0, 36.9, 32.5, 29.2, 27.0, 25.8, 24.2 (2 quaternary ar. C not detected). HRMS (ESI^+^) *m/z* [M+H]^+^ 399.1810; calculated for C_19_H_23_N_6_O_4_: 399.1781.

*Synthesis of **9**:* Lenalidomide (500 mg, 1.93 mmol, 1.0 eq) and hex-5-ynoic acid (0.320 mL, 2.89 mmol, 1.5 eq) were dissolved in dry DMF (16 mL) stirring under nitrogen atmosphere. HATU (734 mg, 1.93 mmol, 1.0 eq) and DIPEA (1.0 mL, 5.79 mmol, 3.0 eq) were added and the mixture was stirred overnight at room temperature (TLC monitoring: DCM/MeOH 9:1). Then, water and sat. aqueous solution of NH_4_Cl were added, and the solution was extracted with EtOAc (5 x 20 mL). The combined organic layers were dried over Na_2_SO_4_, filtered, and concentrated under reduced pressure. The crude was purified by flash column chromatography (eluent mixture: DCM/MeOH 92:8) to obtain **9** (526 mg, 1.49 mmol, 77% yield) as a white solid. ^1^H NMR (400 MHz, pyridine-*d_5_*) δ 12.86 (s, 1H), 10.78 (s, 1H), 8.00 (d, *J* = 7.8 Hz, 1H), 7.88 (d, *J* = 7.8 Hz, 1H), 7.47 (t, *J* = 7.8 Hz, 1H), 5.66 (dd, *J* = 13.4, 5.0 Hz, 1H), 4.84-4.67 (m, 2H), 2.96 (ddd, *J* = 18.2, 13.3, 5.3 Hz, 1H), 2.86-2.82 (m, 1H), 2.80-2.77 (m, 2H), 2.69 (s, 1H), 2.40-2.34 (m, 3H), 2.12-2.05 (m, 3H). ^13^C NMR (101 MHz, pyridine-*d_5_*) δ 173.8, 172.2, 171.6, 169.7, 135.6, 135.4, 134.5, 129.6, 126.5, 120.8, 84.9, 71.4, 53.1, 48.2, 35.9, 32.7, 25.2, 24.3, 18.8. HRMS (ESI^+^) *m/z* [M+H]^+^ 354.1442; calculated for C_19_H_20_N_3_O_4_: 354.1454.

*Synthesis of **10***: A mixture of 6-heptynoic acid (18 µL, 0.14 mmol, 1.2 eq), HATU (88 mg, 0.24 mmol, 2.0 eq) and DIPEA (60 µL, 0.35 mmol, 3.0 eq) in DCM (1.2 mL) was stirred for 30 mintutes at room temperature, and then VHL ligand (50 mg, 0.12 mmol, 1.0 eq) was added. The reaction was stirred at room temperature overnight and monitored by TLC (eluent: DCM/MeOH 95:5). After completion, water was added, and the mixture was extracted with EtOAc (3 x 10 mL). The combined organic layer was washed with brine (1 x 10 mL), dried over anhydrous Na_2_SO_4_, filtered, and concentrated under reduced pressure. The crude was purified by flash column chromatography (eluent: DCM/MeOH 97:3) affording compound **10** (28 mg, 0.052 mmol, 45% yield) as a white powder. ^1^H NMR (400 MHz, CDCl_3_) δ 8.72 (s, 1H), 7.38 – 7.33 (m, 4H), 7.29 – 7.26 (m, 1H), 6.29 (d, *J* = 8.6 Hz, 1H), 4.71 (t, *J* = 7.5 Hz, 1H), 4.61 – 4.51 (m, 3H), 4.33 (dd, *J* = 15.0, 5.2 Hz, 1H), 4.11 (d, *J* = 11.5, 1H), 3.61 (dd, *J* = 11.5, 3.5 Hz, 1H), 2.52 (s, 3H), 2.36 (t, *J* = 7.5 Hz, 1H), 2.25 – 2.17 (m, 4H), 2.16 – 2.10 (m, 1H), 1.94 (t, *J* = 2.7 Hz, 1H), 1.76 - 1.68 (m, 2H), 1.58 – 1.49 (m, 2H), 0.93 (s, 9H). HRMS (ESI^+^) *m/z* [M+H]^+^ 539.2671; calculated for C_29_H_39_N_4_O_4_S: 539.2692. Spectroscopic data are consistent with that reported in the literature^35^.

*Synthesis of **1***: N_3_-maytansinoid **5** (30 mg, 0.041 mmol, 1 eq) and alkyne-thalidomide derivative **6** (24 mg, 0.061 mmol, 1.5 eq) were dissolved in DMSO (0.96 mL). Then, water (0.16 mL), CuSO_4_·5H_2_O (16 mg,0.065 mmol, 1.6 eq) and sodium ascorbate (48 mg, 0.24 mmol, 6 eq) were added and the mixture was stirred at room temperature for 24 h, monitoring by TLC (eluent: DCM/MeOH 95:5). Water was added and the aqueous layer was extracted in DCM (3 x 5 mL). The organic layer was washed with H_2_O (1 x 5 mL) and brine (1 x 5 mL), dried over Na_2_SO_4_ and concentrated in vacuum. The crude was purified *via* flash chromatography (eluent: DCM/MeOH from 98:2 to 96:4) to provide product **1** (17 mg, 0.015 mmol, 37% yield) as a yellow powder. ^1^H NMR (400 MHz, MeOD) δ 7.95 (d, *J* = 8.0 Hz, 2H), 7.71 (d, *J* = 1.8 Hz, 1H), 7.53 (t, *J* = 7.6 Hz, 1H), 7.33 – 7.29 (m, 2H), 7.15 (d, *J* = 1.8 Hz, 1H), 7.09 – 7.02 (m, 3H), 6.47 (dd, *J* = 15.4, 11.0 Hz, 1H), 5.90 (d, *J* = 11.0 Hz, 1H), 5.03 (dd, *J* = 12.4, 5.4 Hz, 1H), 4.96 (t, *J* = 5.4 Hz, 1H), 4.91 – 4.85 (m, 1H), 4.72 – 4.67 (m, 2H), 4.60 – 4.52 (m, 2H), 4.23 – 4.10 (m, 1H), 3.99 (s, 3H), 3.68 (m, 2H), 3.65 – 3.57 (m, 4H), 3.51 – 3.46 (m, 3H), 3.41 (d, *J* = 9.0, 1H), 3.34 – 3.32 (m, 2H), 3.30 – 3.38 (m, 1H), 3.25 (s, 3H), 3.13 (s, 3H), 3.01 – 2.98 (m, 1H), 2.84 – 2.68 (m, 3H), 2.28 (dd, *J* = 14.2, 3.2 Hz, 1H), 2.19 – 2.04 (m, 2H), 1.70 (s, 3H), 1.55 – 1.42 (m, 3H), 1.23 (d, *J* = 6.0 Hz, 3H), 0.87 (s, 3H). ^13^C NMR (101 MHz, MeOD) δ 174.6, 172.9, 171.6, 171.2, 170.7, 169.3, 167.3, 157.7, 155.3, 148.3, 145.9, 144.9, 143.2, 142.9, 141.1, 137.6, 137.2, 133.7, 131.5, 129.9, 129.7, 129.3, 128.8, 125.9, 125.3, 122.8, 119.9, 118.4, 114.9, 112.0, 111.3, 90.1, 81.6, 78.7, 75.8, 71.5, 70.73, 70.70, 68.2, 67.5, 65.1, 57.3, 57.0, 52.2, 50.3, 47.6, 43.3, 39.8, 37.3, 36.1, 33.9, 32.2, 30.2, 23.8, 15.9, 14.8, 13.0. HRMS (ESI^+^) *m/z* [M+Na]^+^ 1159.4143; calculated for C_57_H_65_ClN_8_O_15_Na: 1159.4156.

*Synthesis of **2***: N_3_-lenalidomide derivative **8** (20 mg, 0.050 mmol, 1.5 eq) and maytansinoid **7** (23 mg, 0.033 mmol, 1.0 eq) were dissolved in DMSO (0.72 mL). Then, water (0.12 mL), CuSO_4_·5H_2_O (13 mg,0.053 mmol, 1.6 eq) and sodium ascorbate (40 mg, 0.20 mmol, 6.0 eq) were added and the mixture was stirred at room temperature for 24 h, monitoring by TLC (eluent: DCM/MeOH 95:5). Water was added and the aqueous layer was extracted in DCM (3 x 5 mL). The organic layer was washed with H_2_O (1 x 5 mL) and brine (1 x 5 mL), dried over Na_2_SO_4_ and concentrated in vacuum. The crude was purified *via* flash chromatography (eluent: DCM/MeOH from 98:2 to 95:5) to provide product **2** (23 mg, 0.021 mmol, 65% yield) as a white powder. ^1^H NMR (400 MHz, MeOD) δ 7.77 (s, 1H), 7.73-7.70 (m, 1H), 7.63 (dt, *J* = 7.6, 1.0 Hz, 1H), 7.50 (t, *J* = 7.6 Hz, 1H), 7.13 (d, *J* = 1.8 Hz, 1H), 6.85 (d, *J* = 1.8 Hz, 1H), 6.61 (dd, *J* = 15.4, 11.0 Hz, 1H), 6.26 (d, *J* = 11.0 Hz, 1H), 5.55 (dd, *J* = 15.4, 8.9 Hz, 1H), 5.16 (dd, *J* = 13.2, 5.0 Hz, 1H), 4.77 (dd, *J* = 12.0, 3.0 Hz, 1H), 4.48 (s, 2H), 4.37 (t, *J* = 7.5 Hz, 2H), 4.17 (td, *J* = 10.9, 3.0 Hz, 1H), 3.97 (s, 3H), 3.58 – 3.56 (m, 2H), 3.35 (d, *J* = 1.5 Hz, 3H), 3.31 – 3.28 (m, 1H), 3.05 (s, 3H), 2.95-2.86 (m, 1H), 2.81 - 2.72 (m, 4H), 2.64 - 2.48 (m, 4H), 2.43 (t, *J* = 7.5 Hz, 2H), 2.21-2.17 (m, 1H), 2.11 (dd, *J* = 13.5, 3.0 Hz, 1H), 1.92 (p, *J* = 7.5 Hz, 2H), 1.77 - 1.67 (m, 9H), 1.58 - 1.47 (m, 3H), 1.37 (p, *J* = 7.5 Hz, 2H), 1.19 (d, *J* = 6.3 Hz, 3H), 0.86 (s, 3H). ^13^C NMR (101 MHz, MeOD) δ 174.6, 174.2, 173.8, 172.1, 171.3, 171.0, 157.5, 155.2, 148.7, 143.0, 142.8, 141.1, 136.3, 134.6, 133.9, 133.8, 130.1, 129.8, 127.8, 126.0, 123.4, 123.1, 121.4, 119.7, 115.0, 89.7, 81.9, 77.9, 75.9, 67.9, 61.9, 57.2, 57.0, 53.7, 51.0, 47.5, 39.2, 37.5, 37.0, 36.2, 34.8, 33.8, 32.4, 31.0, 30.7, 29.8, 27.0, 26.1, 26.0, 25.4, 24.1, 15.9, 14.7, 12.5. HRMS (ESI^+^) *m/z* [M+H]^+^ 1071.4602; calculated MS for C_54_H_68_ClN_8_O_13_: 1071.4594.

*Synthesis of **3***: N_3_-maytansinoid **5** (30 mg, 0.041 mmol, 1.0 eq) and alkyne-lenalidomide derivative **9** (22 mg, 0.061 mmol, 1.5 eq) were dissolved in DMSO (0.96 mL). Then, water (0.16 mL), CuSO_4_·5H_2_O (16 mg,0.065 mmol, 1.6 eq) and sodium ascorbate (48 mg, 0.24 mmol, 6.0 eq) were added and the mixture was stirred at room temperature for 24 h, monitoring by TLC (eluent: DCM/MeOH 95:5). Water was added and the aqueous layer was extracted in DCM (3 x 5 mL). The organic layer was washed with H_2_O (1 x 5 mL) and brine (1 x 5 mL), dried over Na_2_SO_4_ and concentrated in vacuum. The crude was purified *via* flash chromatography (eluent: DCM/MeOH from 98:2 to 95:5) to provide product **3** (21 mg, 0.019 mmol, 46% yield) as a white powder. ^1^H NMR (400 MHz, acetone-d_6_) δ 9.80 (s, 1H), 9.24 (s, 1H), 8.00-7.98 (m, 3H), 7.61 (d, *J* = 1.1 Hz, 1H), 7.54 (dd, *J* = 7.5, 1.1 Hz, 1H), 7.48 (t, *J* = 7.5 Hz, 1H), 7.41-7.39 (m, 2H), 7.23 (d, *J* = 2.0 Hz, 1H), 7.12 (d, *J* = 2.0 Hz, 1H), 6.58 (dd, *J* = 15.4, 11.0 Hz, 1H), 6.40 – 6.37 (m, 1H), 6.06 (dd, *J* = 11.0, 5.6 Hz, 1H), 5.20 (ddd, *J* = 13.3, 5.1, 1.3 Hz, 1H), 5.02-4.91 (m, 2H, H3), 4.78 – 4.63 (m, 2H), 4.55 – 4.48 (m, 2H), 4.19 – 4.13 (m, 1H), 4.01 (d, *J* = 1.2 Hz, 3H), 3.58 (d, *J* = 12.4 Hz, 1H), 3.47 (d, *J* = 9.0 Hz, 1H), 3.36 – 3.31 (m, 3H), 3.23 (d, *J* = 2.0 Hz, 3H), 3.09 (d, *J* = 1.5 Hz, 3H), 3.00 – 2.94 (m, 2H), 2.82 – 2.79 (m, 2H), 2.71 (t, *J* = 7.3 Hz, 2H), 2.57 – 2.48 (m, 1H), 2.47 – 2.39 (m, 2H), 2.25 – 2.19 (m, 2H), 2.01 - 1.97 (m, 2H), 1.74 (s, 3H), 1.55 (d, *J* = 14.0 Hz, 1H), 1.49 – 1.41 (m, 2H), 1.17 (dd, *J* = 6.4, 2.5 Hz, 3H), 0.93 (s, 3H). ^13^C NMR (101 MHz, acetone-d_6_) δ 172.82, 172.80, 172.1, 171.4, 169.0, 166.5, 157.1, 152.2, 152.1, 147.5, 144.60, 144.58, 143.2, 142.3, 140.4, 135.10, 135.08, 134.1, 133.1, 131.0, 129.7, 129.5, 125.7, 125.6, 122.93, 122.91, 122.8, 120.0, 119.5, 114.5, 89.8, 81.5, 78.2, 74.6, 67.2, 61.4, 57.1, 56.8, 52.84, 52.82, 51.5, 47.3, 47.2, 39.5, 37.2, 36.4, 35.5, 33.6, 32.2, 26.18, 26.15, 25.4, 24.0, 15.8, 14.9, 13.0. HRMS (ESI^+^) *m/z* [M+Na]^+^ 1113.4131; calculated for C_56_H_63_ClN_8_O_13_Na: 1113.4101.

*Synthesis of **4***: N_3_-maytansinoid **5** (26 mg, 0.036 mmol, 1.0 eq) and alkyne-VHL ligand derivative **10** (25 mg, 0.046 mmol, 1.3 eq) were dissolved in DMSO (0.77 mL). Then, water (0.13 mL), CuSO_4_·5H_2_O (14 mg,0.057 mmol, 1.6 eq) and sodium ascorbate (42 mg, 0.21 mmol, 6.0 eq) were added and the mixture was stirred at room temperature for 24 h, monitoring by TLC (eluent: DCM/MeOH 95:5). Water was added and the aqueous layer was extracted in DCM (3 x 5 mL). The organic layer was washed with H_2_O (1 x 5 mL) and brine (1 x 5 mL), dried over Na_2_SO_4_ and concentrated in vacuum. The crude was purified *via* flash chromatography (eluent: DCM/MeOH from 98:2 to 96:4) to provide the product **4** (19 mg, 0.015 mmol, 41% yield) as a pale-yellow powder. ^1^H NMR (400 MHz, acetone-d_6_) δ 8.85 (s, 1H), 7.99 (d, *J* = 8.2 Hz, 2H), 7.78 (m, 1H), 7.55 (s, 1H), 7.49 – 7.37 (m, 6H), 7.25 (d, *J* = 1.8 Hz, 1H), 7.14 (d, *J* = 1.8 Hz, 1H), 6.58 (dd, *J* = 15.5, 11.0 Hz, 1H), 6.42 (s, 1H), 6.04 (d, *J* = 11.0 Hz, 1H), 5.00 (dd, *J* = 15.5, 9.2 Hz, 1H), 4.95 (dd, *J* = 12.5, 3.6 Hz, 1H), 4.73 – 4.57 (m, 6H), 4.35 (dd, *J* = 15.5, 5.1 Hz, 1H), 4.20 (ddd, *J* = 12.5, 10.6, 2.2 Hz, 1H), 4.02 (s, 3H), 3.95 – 3.92 (m, 1H), 3.75 (dd, *J* = 10.6, 4.1 Hz, 1H), 3.58 (d, *J* = 12.5 Hz, 1H), 3.48 (d, *J* = 9.2 Hz, 1H), 3.36 – 3.32 (m, 3H), 3.24 (s, 3H), 3.10 (s, 3H), 3.01 (d, *J* = 9.6 Hz, 1H), 2.84 – 2.78 (m, 1H), 2.63 – 2.59 (m, 2H), 2.47 (s, 3H), 2.36 – 2.11 (m, 5H), 1.75 (s, 3H), 1.68 – 1.53 (m, 5H), 1.52 – 1.38 (m, 2H), 1.21 (d, *J* = 6.5 Hz, 3H), 0.99 (s, 9H), 0.94 (s, 3H).^13^C NMR (101 MHz, acetone-d_6_) δ 173.4, 172.6, 171.7, 169.0, 166.6, 157.1, 152.2, 151.3, 149.2, 148.0, 144.7, 143.3, 142.3, 140.5, 140.3, 133.1, 131.4, 131.0, 130.0, 129.9, 129.8, 129.7, 129.2, 128.7, 125.8, 123.0, 122.4, 119.6, 114.5, 89.9, 81.5, 78.3, 74.7, 70.7, 67.3, 61.5, 60.0, 57.9, 57.5, 57.1, 56.8, 51.4, 47.2, 43.2, 39.6, 38.3, 37.30, 37.28, 36.13, 36.07, 35.5, 33.6, 27.0, 25.9, 25.8, 16.3, 15.8, 14.9, 13.0. HRMS (ESI^+^) m/z [M+Na]^+^ 1298.5342; calculated for C_66_H_82_ClN_9_O_13_S: 1298.5339.

### Protein expression and purification

Lyophylized calf brain tubulin was bought from the group of J. F. Díaz (Centro de Investigaciones Biológicas Margarita Salas, CSIC, Madrid). Purified DDB1τιB-CRBN-complex was a gift from the group of N. Thomä (Friedrich Miescher Institute for Biomedical Research, Basel and EPFL, Ecublens).

The VCB-complex was expressed and purified using an adapted protocol from Gadd and co-workers.^36^ Briefly, N-terminally His6-tagged VHL (54–213), Elongin C (17–112) and Elongin B (1–104) were co-expressed (VHL and EloB/EloC plasmids provided by Prof. Ciulli, University of Dundee, UK) in *Escherichia coli* BL21(DE3) at 20 °C for 16 h using 0.5 mM isopropyl β-D-1-thiogalactopyranoside (IPTG). *E. coli* cells were harvested by centrifugation and lysed with an ultrasonic cell disruptor and lysate clarified by centrifugation for 30 minutes at 4000 rpm at 4 °C. His6-tagged VCB was purified on a IMAC HisTrap column by elution with an imidazole gradient (20 mM to 500 mM). The His6 tag was cleaved overnight using TEV protease during dialysis into low-concentration imidazole buffer. VCB was then passed through the HisTrap column a second time and the flow-through was collected. VCB was then additionally purified by anion exchange and size-exclusion chromatography using XK 16/20 and Superdex-75 16/60 columns, respectively. The final purified complex was concentrated to 20 mg/mL and stored at -80 °C in 20 mM HEPES, pH 7.5, supplemented with 100 mM sodium chloride and 1 mM β-mercaptoethanol.

### Biophysical characterization

#### Nanoscale Differential Scanning Fluorimetry (nanoDSF)

Real-time simultaneous monitoring of the ITF (Intrinsic Tryptophan Fluorescence) at 330 nm and 350 nm for tubulin/CRBN/VCB alone and in complex with compounds was carried out in a Prometheus Panta instrument (NanoTemper Technologies) with an excitation wavelength of 280 nm. Before use, tubulin was resuspended in BRB80 buffer (80 mM PIPES/KOH, pH 6.8, supplemented with 1 mM MgCl_2_ and 1 mM EGTA) over 20 minutes on ice and then ultracentrifuged to remove aggregates at 50’000 rpm using a TLA120.1 rotor for 10 min at 4°C. The concentration of tubulin was measured using absorbance at 280 nM (Nanodrop One, ε_tubulin_=115 000 cm^-1^M^-1^) and tubulin diluted into MES buffer (50 mM pH 5.5) to a final concentration of 3 μM (0.3 mg/mL). Compounds were added to a 10x excess and DMSO was added in equal amounts to the tubulin control. Capillaries were filled with 10 μl of sample, placed into the sample holder and the temperature was increased from 20/25 to 90 °C at a ramping rate of 1 °C/min. The ratio of the recorded emission intensities (Em350 nm/Em330 nm), which represents the change in Trp fluorescence intensity was plotted as a function of the temperature. The fluorescence intensity ratio, its first derivative and the plots (Supplementary Figures S1-S3) were obtained using the manufacturer’s software (PR.ThermControl, version 1.2). Two independent measurements were carried out for each condition.

#### Size Exclusion Chromatography coupled to Multiangle Light Scattering (SEC-MALS)

Lyophilized tubulin was resuspended in BRB80 (Pipes/KOH pH 6.8 80 mM, MgCl_2_ 1 mM, EGTA 1 mM), and incubated on ice for 10 min. Then samples were ultracentrifuged for 10 min at 50’000 rpm using a TLA120.1 rotor. The tubulin concentration was measured using absorbance at 280 nm (Nanodrop One, ε_tubulin_=115 000 cm^-1^M^-1^). Samples containing tubulin (10 μM), CRBN (10 μM), VCB (10 μM) as references, as well as mixtures of tubulin and each ligase were prepared. Additionally, samples of tubulin (10 μM) and CRBN (10 μM) containing PROTACs **1-3** (15 μM) and tubulin (10 μM) and VCB (10 μM) with **4** (24 μM) were prepared. All samples were equilibrated for 1 h on ice, before SEC-MALS measurements were performed at 25 °C. A GE Healthcare Superdex 200 Increase 10/300 SEC column on an Agilent 1260 HPLC at a flowrate of 0.5 mL/min of BRB80 running buffer was used. The column was equilibrated overnight with running buffer to obtain stable baseline signals from the detectors before data collection. Absorbance at 280 nm was monitored with an Agilent multi-wavelength absorbance detector to monitor protein elution. The signals for multiangle light scattering and differential refractive index were measured using a Wyatt Heleos II 8+ detector and a Wyatt Optilab detector, respectively. Obtained chromatograms were corrected for inter-detector delay volumes and band broadening, and the signals of light-scattering detector normalized by calibration with 2 mg/mL injections of BSA solutions (Thermo Pierce) using standard protocols in ASTRA 8.1.2. The weight-averaged molar mass (Mw) of the samples was calculated using the ASTRA 8.1.2 software (Wyatt Technology).

### X-ray crystallography structure determination

T_2_R-TTL crystals were prepared according to the previously established protocols^28, 37^. Crystals were grown at room temperature over a week in a buffer solution containing PEG 4K (4 %), glycerol (13 or 14 %), MgCl_2_ (30 mM), CaCl_2_ (30 mM), MES/imidazole pH 6.5 (100 mM) and tyrosine (5 mM). Compounds **1-4** were soaked to the crystals at a concentration of 2.5 or 5 mM for 6 hours. X-ray diffraction data at 100 K and 1 Å wavelength were collected at beamline X06SA of the Swiss Light Source, Paul Scherrer Institut, 5332 Villigen PSI, Switzerland. Data were processed using XDS^38^ and the structures were determined by the difference Fourier method based on the phases of a T_2_R-TTL model in the absence of ligands and solvent molecules (PBD ID: 5LXT). Refinements were done in PHENIX^39^ and model building in Coot^40^. Ligand restraints were generated with eLBOW (part of the PHENIX suite) upon energy minimization of the individual ligands in Moloc^41^. The structures of the T_2_R-TTL-**1**, **2**, **3**, and **4** complexes were refined to 2.2-2.3 Å resolution. Detailed data collection and refinement statistics are listed in the Supporting Information (Table S4). The coordinates and structure factors were deposited at the Protein Data Bank (RCSB PDB, www.rcsb.org) under accession numbers PDB: 9H34 (T_2_R-TTL-**1**), 9H33 (T_2_R-TTL-**2**), 9H32 (T_2_R-TTL-**3**) and 9H31 (T_2_R-TTL-**4**). Molecular graphics were created using PyMOL, The PyMOL Molecular Graphics System, Version 2.3.4 Schrödinger, LLC.

Crystals of the VCB-**4** complex were grown at 4 °C by the hanging-drop vapor-diffusion method by mixing 0.2 μL of VCB complex at 15 mg/mL preincubated with 1 mM **4** with 0.2 μL reservoir solution. Crystals grew after 4-7 days incubation in the presence of 9-11% PEG 8000, 0.2 M magnesium acetate and 0.1 M sodium cacodylate at pH 5.7^42^. Crystals were flash-cooled in liquid nitrogen following subsequent transfer into two cryo-protectant solutions supplemented with 18% and 22% PEG2000. X-ray diffraction data were collected at 100 K and 1 Å wavelength at beamline X06SA of the Swiss Light Source, Paul Scherrer Institut, 5332 Villigen PSI, Switzerland. Data were processed using XDS^38^, and the structure was solved by molecular replacement using the VCB structure (PDBID 6HAY, chains B, C and D) as the template model, with ligands and solvents removed. The phases were determined by using CCP4-8.0-Phaser^43^. Refinement was carried in PHENIX^39^, while model building was done in Coot^40^. The structure of VCB**-4** was refined to 2.5 Å resolution. Detailed data collection and refinement statistics are provided in the Supporting Information (Table S4). The coordinates and structure factors were deposited at the Protein Data Bank (RCSB PDB, www.rcsb.org) under accession code 9H30 (VCB-**4**). Molecular graphics were created using PyMOL, The PyMOL Molecular Graphics System, Version 2.3.4 Schrödinger, LLC.

### Cell biology

#### Cell culture and treatments

Caco-2 cells were cultured in DMEM supplemented with 10% FBS, 2mM L-Glutamine, and 1% Penicillin/Streptomycin antibiotic. One day prior to the harvesting, cells were plated at maximum confluence (1.25×10^5^ cells/cm^2^). The next day, the compounds were administered for 4 hours at different concentrations, and 0.1% DMSO (v/v) was used as vehicle control.

#### Cytotoxicity assays

Hoechst test was performed to assess compounds cytotoxicity, according to manufacturer instructions. CaCo-2 cells were incubated with increasing concentrations of each compound, 0.1% DMSO was used as control. After 48 hours, fluorescence at 461 nm was read at an Ensight® multimode plate reader (Perkin Elmer).

#### Western blot

Total cellular extracts were collected by scraping under Laemmli buffer incubation (2% SDS, 10% glycerol, 5% β-mercaptoethanol, 0.001% bromophenol blue, and 62.5 mmol/L Tris, pH 6.8) supplemented with protease inhibitors cocktail 1:1000 (M250, AMRESCO). Protein concentration was measured with Micro BCA TM Protein Assay Kit (Thermo Scientific). The electrophoretic run was carried out on Bis-Tris polyacrylamide gels, then proteins were transferred to methanol-preactivated PVDF membrane (Immobilon-F transfer membrane, Millipore) under constant voltage (60V) for 90 minutes at 4°C. A blocking step was done by incubating membranes with a solution of 5% BSA in Tris-buffered saline (TBS), 0.1% Tween, for 1 hour at room temperature. Then, membranes were incubated ON at 4°C with the following primary antibodies: anti α-Tubulin mouse IgG antibody (1:2000; T6074 Sigma); anti actin rabbit IgG antibody (1:2000; A2066, Sigma). The incubation with secondary antibodies was performed for 1 hour at room temperature, in the dark, using either the Alexa Fluor 488 donkey anti rabbit antibody (1:4000; Invitrogen, A21206), or the Alexa Fluor 555 goat anti mouse antibody (1:4000; Invitrogen, A32727) diluted in in BSA 1% in TBS, 0.1% Tween. Images were acquired at Chemidoc Touch (Biorad) and relative band intensity levels were quantified with ImageLab software (Biorad).

## Author contributions

A. Maiocchi: design of experiments, chemical synthesis, crystallographic structure determination and analysis, biochemistry work and data analysis, wrote the first draft of the manuscript and prepared the figures (equal contribution); A-C. Abel: design of experiments, crystallographic structure determination and analysis, biochemistry work and data analysis, wrote the first draft of the manuscript and prepared the figures (equal contribution); M. Baselini: cell biology data acquisition and analysis, figure and manuscript preparation; H. Pérez-Peña: molecular modelling, figure and manuscript preparation; Z. Boiarska: tubulin crystallization and X-ray data collection; E.E. Ferrandi: organic chemistry support, data analysis; Z. Kozicka: recombinant protein expression and purification; V. Fasano: organic chemistry support, data analysis; S. Pieraccini: supervision of molecular modelling, data analysis; G. Cappelletti: supervision of cell biology, data analysis; M.O. Steinmetz: supervision of study, data analysis; A.E. Prota: research design, supervision of study, crystallography, structure determination and analysis, analysis of biophysical data, project coordination, manuscript preparation; D. Passarella: research design, supervision of study, chemistry and data analysis, project coordination, manuscript preparation; All authors discussed the results and commented on the manuscript.

## Acknowledgements

The authors thank Professor Dr. A. Ciulli from University of Dundee (Centre for Targeted Protein Degradation) for providing VHL and EloB/EloC plasmids. Many thanks to Professor Dr. Nicolas Thomä from the Friedrich Miescher Institute for Biomedical Research (Basel) and EPFL (Ecublens) for the supply of the CRBN-DDB1τιB-complex. The authors thank the staff of the Swiss Light Source responsible for beamline X06SA for their excellent support during X-ray data collection. The authors sincerely thank Dr. Tobias Mühlethaler and Dr. Timothy Sharpe (Biozentrum of the University of Basel) for their support in carrying out the SEC-MALS experiments in their facility. This work was partially supported by the H2020-MSCA-ITN-2019 (860070 TUBINTRAIN to S.P., G.C., A.E.P, and D.P.) and the Swiss National Science Foundation (grant 310030_192566; to M.O.S.).

## Data Availability Statement and competing interest

The data that support the findings of this study are available in the supplementary material of this article. The coordinates and structure factors of the T_2_R-TTL-PROTAC-**1-4** and the VHL-ElonginC-ElonginB-PROTAC-**4** complexes have been deposited at the Protein Data Bank (RCSB PDB, https://www.rcsb.org/) under the accession numbers 9H34, 9H33, 9H32,9H31 and 9H30, respectively. The authors declare no competing interests.

