## Supplemental Information for "Effective Tubulin Degradation by Rationally Designed Proteolysis Targeting Chimeras"

### Index

#### Chemical functionalization of maytansinol POI ligand: synthesis of moieties 5 and 7

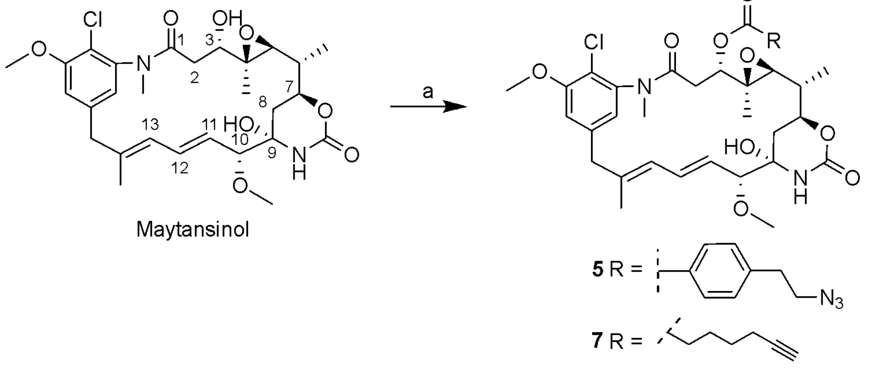

**Supplementary Scheme 1**: Synthesis of target C3-acylated maytansinol derivatives **5** and **7**. a) 6-heptinoic acid or 4-(2-azidoethyl)benzoic acid, DMAP, DCC, DCM, rt, 5 h, **5**: 45% yield, **7**: 35% yield.

#### Chemical functionalization of E3 binders: synthesis of alkynyl and azido derivatives 6-10

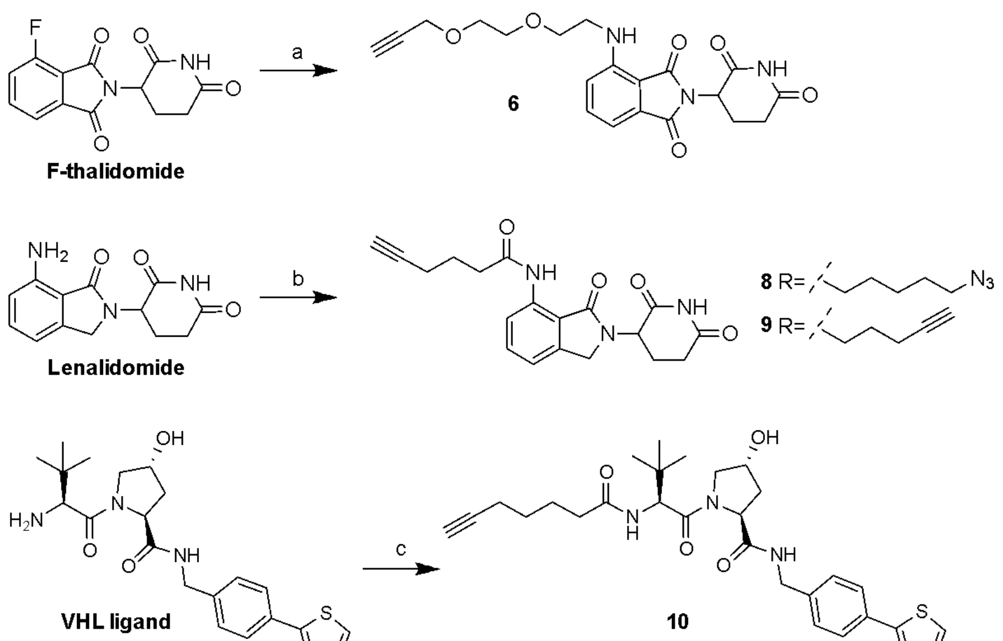

**Supplementary Scheme 2**: Synthesis of functionalized E3-targeting moieties **6-10**. a) 2-(2-(prop-2-yn-1-yloxy)ethoxy)ethan-1-amine, DIPEA, DMF, 90 °C, 5 h, 49% yield; b) 6-azidohexanoic acid or 5-hexynoic acid, HATU, DIPEA, DMF, rt, 24 h, **8**: 48% yield, **9**: 77% yield; c) 6-heptynoic acid, HATU, DIPEA, DCM, rt, 24 h, 45% yield.

#### Thermal Shift Assays for binary binding (NanoDSF)

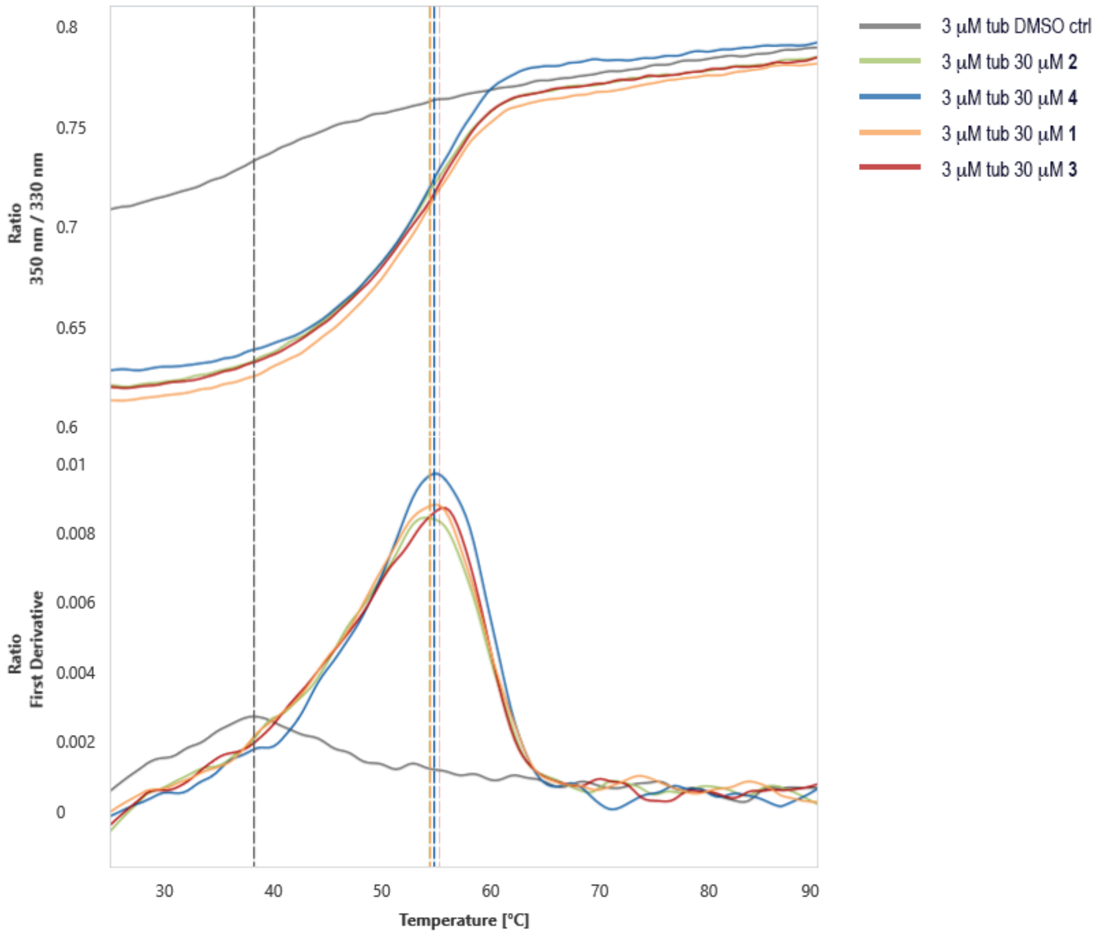

**Supplementary Fig. S1:** Thermal shift assay binary binding to tubulin. Representative nanoDSF experiments for measurements of the melting points of 3 µM tubulin samples in the absence and presence of 30 µM PROTACs **1-4** in 50 mM MES buffer pH 5.5 are shown. The ratio of fluorescence emission at 350/330 nm wavelengths and the first derivative thereof are plotted against the temperature, detected melting points are indicated by vertical dashed lines colour-coded according to the sample.

| **Sample** | **Buffer** | **°C** |
|---|---|---|
| tubulin 3 µM | MES pH 5.5 | 39.6 ± 1.9 |
| tubulin 3 µM + **4** 30 µM | MES pH 5.5 | 55.1 ± 0.3 |
| tubulin 3 µM + **2** 30 µM | MES pH 5.5 | 54.6 ± 0.6 |
| tubulin 3 µM + **3** 30 µM | MES pH 5.5 | 55.2 ± 0.1 |
| tubulin 3 µM + **1** 30 µM | MES pH 5.5 | 54.6 ± 0.2 |
| not shown: tubulin 3 µM | BRB80 | 59.0 ± 0.1 |

**Supplementary Table S1**: Melting points measured in the above shown nanoDSF experiments for samples of 3 µM tubulin in absence and presence of 30 µM PROTACs **1-4** in MES 50 mM pH 5.5 buffer and as reference in BRB80 buffer (not shown in the Figure S1). Values are the mean of two technical replicas.

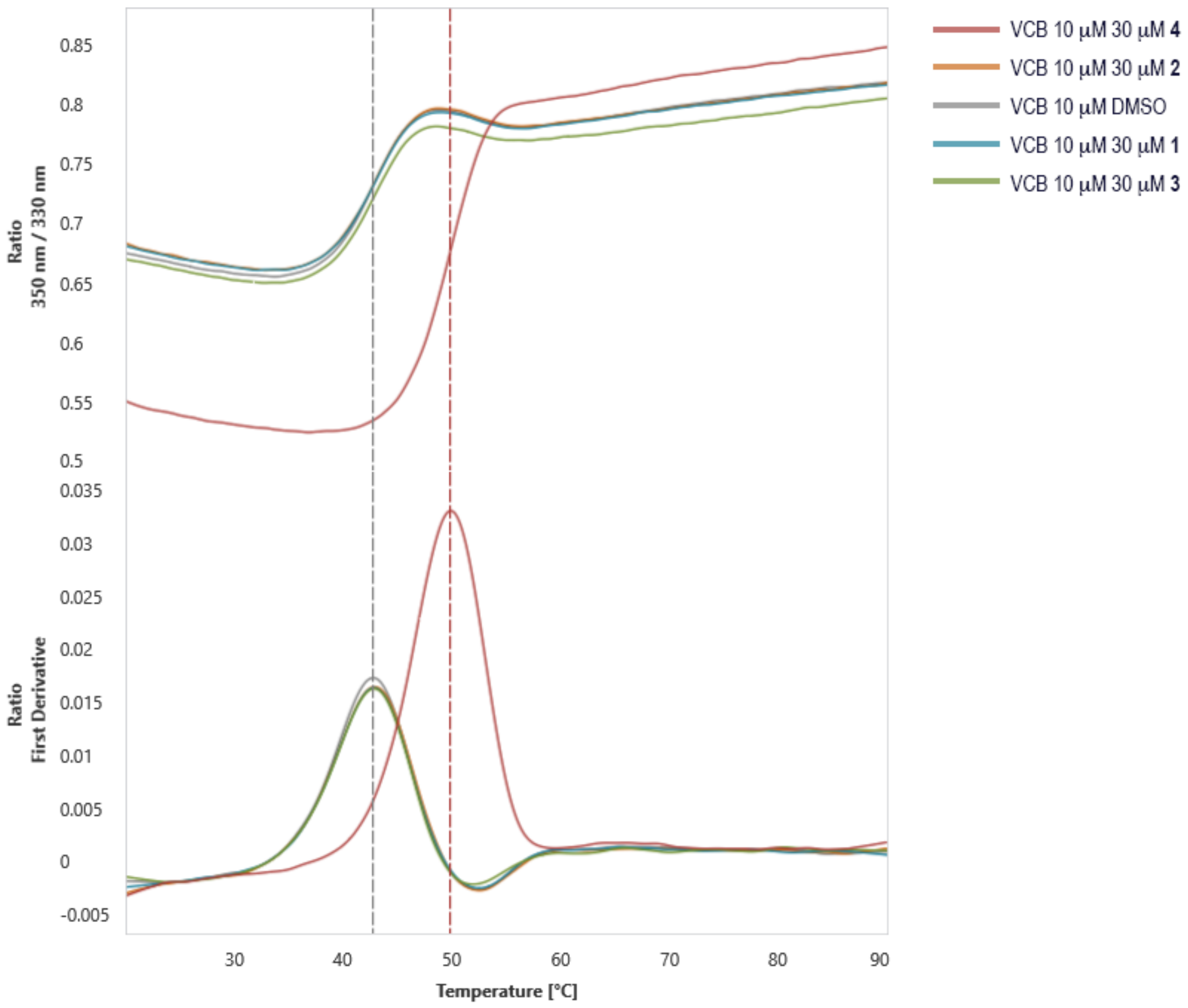

**Supplementary Fig. S2**: Thermal shift assay binary binding to VCB complex. Representative nanoDSF experiments for measurements of the melting points of 10 μM VCB complex samples in the absence and presence of 30 μM PROTACs **1-4** in BRB80 buffer. The ratio of fluorescence emission at 350/330 nm wavelengths and the first derivative thereof are plotted against the temperature, detected melting points are indicated by vertical dashed lines colour-coded according to the sample.

**Supplementary Table S2**: Melting points measured in the above shown nanoDSF data of 10 μM VCB complex in absence and presence of 30 μM PROTACs **1-4** in BRB80 buffer. Values are the mean of two technical replicas.

| **Sample** | **Buffer** | **°C** |
|---|---|---|
| VCB 10 μM | BRB80 | 42.8 ± 0.1 |
| VCB 10 μM + **2** 30 μM | BRB80 | 42.9 ± 0.0 |
| VCB 10 μM + **3** 30 μM | BRB80 | 42.8 ± 0.0 |
| VCB 10 μM + **1** 30 μM | BRB80 | 42.8 ± 0.0 |
| VCB 10 μM + **4** 30 μM | BRB80 | 50.0 ± 0.3 |

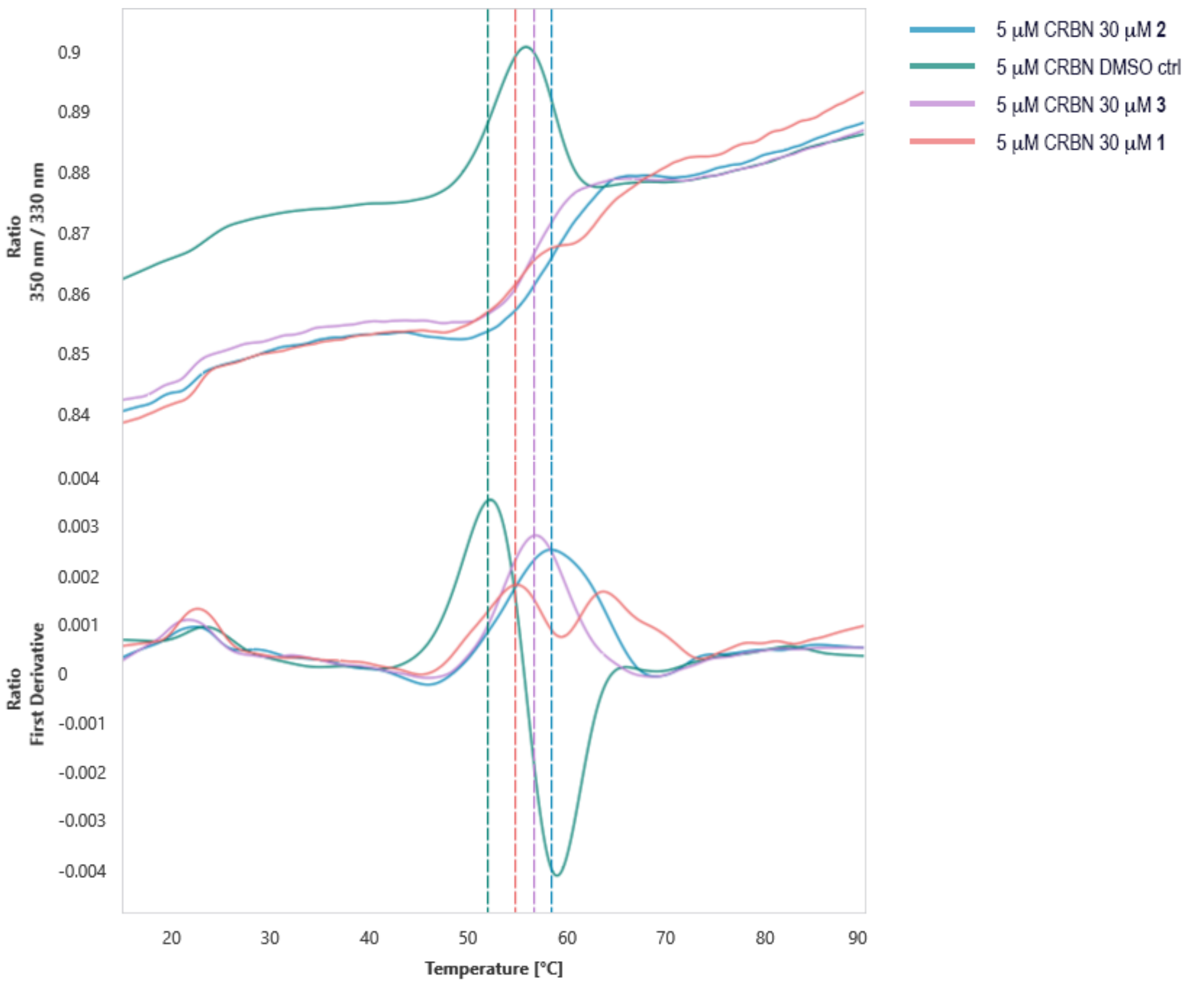

**Supplementary Fig. S3**: Thermal shift assay binary binding to CRBN complex. Representative nanoDSF experiments for measurements of the melting points of 5 µM CRBN complex samples in the absence and presence of 30 µM PROTACs **1-3** in BRB80 buffer. The ratio of fluorescence emission at 350/330 nM wavelengths and the first derivative thereof are plotted against the temperature, detected melting points are indicated by vertical dashed lines colour-coded according to the sample.

**Supplementary Table S3**: Melting points measured in the above shown nanoDSF data of 5 µM CRBN complex in absence and presence of 30 µM PROTACs **1-3** in BRB80 buffer. Values are the mean of two technical replicas.

| **Sample** | **Buffer** | **°C** |
|---|---|---|
| CRBN 5 µM | BRB80 | 52.0 ± 0.1 |
| CRBN 5 µM + **2** 30 µM | BRB80 | 58.1 ± 0.5 |
| CRBN 5 µM + **3** 30 µM | BRB80 | 56.8 ± 0.0 |
| CRBN 5 µM + **1** 30 µM | BRB80 | 55.0 ± 0.3 |

#### X-ray crystallography

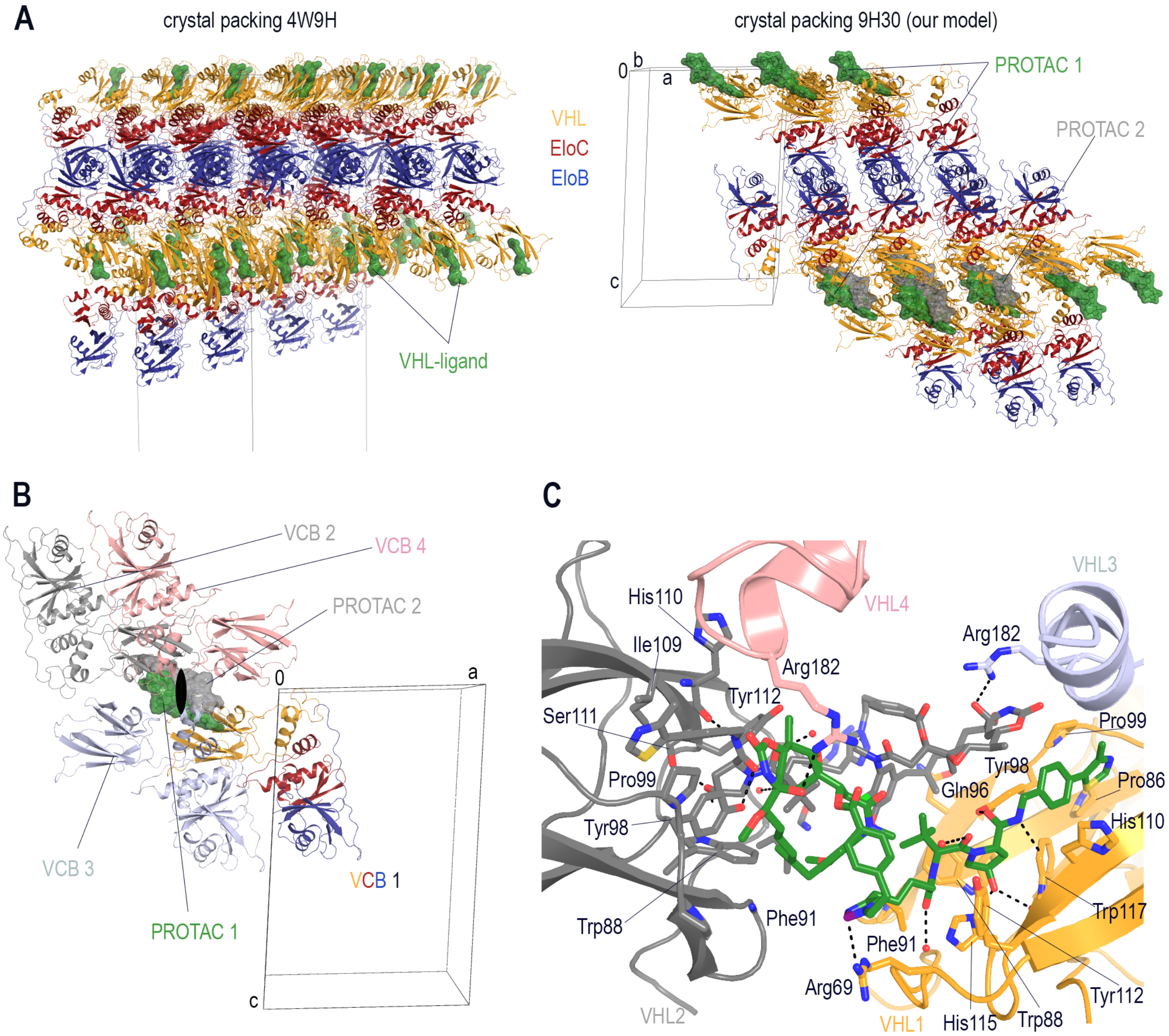

**Supplementary Fig. S4**. VCB crystal packing and coordination of **4** at the interface. (A) Crystal packing of the VCB complexes in the tetragonal crystal form (left, PDB ID 4W9H) compared to the monoclinic crystal form (right, PDB ID 9H30) presented in this study. The VCB complexes are shown in ribbon representation with VHL, Elongin C, and Elongin B coloured yellow, red and blue, respectively. Bound ligands are indicated by green, semi-transparent surface representations, and the unit cells are depicted as black frames. (B) Crystal packing of four adjacent VCB-complexes related by crystallographic symmetry. For the central VCB complex, the same representation as in (A) is applied. The three related VCB complexes are depicted in light blue, grey and light pink with the second PROTAC molecule at the interface shown as grey semi-transparent surface representation. (C) Close-up view of the coordination of two molecules of **4** at the interface of four VHL molecules, coloured as in panel (C) with the two ligand molecules shown as green and grey sticks, respectively. Interacting protein residues are shown as sticks with oxygen, nitrogen and sulphur atoms coloured red, blue and yellow, respectively. Water molecules are depicted as red spheres and hydrogen bond interactions indicated by black dashed lines.

**Description of the interactions of compound 4 at the crystal contact interfaces**

At the crystal contact interface, two molecules of **4** are stacked in a head-to-tail fashion between four VCB complexes, each ligand establishing interactions with three symmetry related protein subunits. The conformation of **4** (bound to VCB 1) is intrinsically stabilised by π-stacking interactions of the triazole linker with the 19-Chloro-20-Methoxy-substituted aromatic ring of the maytansinol core macrocycle that in turn is engaged in hydrophobic interactions with the *tert*-butyl group of the second VH032 moiety bound to a second VCB complex (VCB 2) related by crystallographic symmetry. The conjugated double bonds of the maytansinol scaffold fit into a hydrophobic cleft on the VHL subunit of VCB 2 adjacent to the ligand's binding site. Additionally, the 9-hydroxyl group on the maytansinol scaffold is engaged in a hydrogen bond with Arg182 on the VHL chain of a third symmetry related VCB subunit. Molecule **4** bound to VCB 2 mirrors these interactions, establishing hydrophobic interactions to VCB complex 1 and a hydrogen bond interaction to Arg182 on the fourth symmetry-related VCB complex involved.

**Supplementary Table S4**. X-Ray data collection and refinement statistics.

| | **VCB:4** | **$T_2$R-TTL:4** | **$T_2$R-TTL:1** | **$T_2$R-TTL:2** | **$T_2$R-TTL:3** |
|---|---|---|---|---|---|
| PDB ID | 9H30 | 9H31 | 9H34 | 9H33 | 9H32 |
| *Data collection* | | | | | |
| Wavelength | 0.9999 | 1 | 1 | 1 | 1 |
| Resolution range | 46.78 - 2.5 (2.69 - 2.5) | 49.62 - 2.2 (2.22 - 2.2) | 49.72 - 2.3 (2.33 - 2.3) | 49.61 - 2.3 (2.33 - 2.3) | 48.05 - 2.31 (2.33 - 2.31) |
| Space group | C 1 2 1 | P 21 21 21 | P 21 21 21 | P 21 21 21 | P 21 21 21 |
| Unit cell | 65.5 67.0 101.8 90 94.1 90 | 104.5 156.6 181.3 90 90 90 | 104.4 156.4 181.9 90 90 90 | 105.0 156.7 181.0 90 90 90 | 105.0 157.9 181.4 90 90 90 |
| Total reflections | 104948 (20990) | 18510507 (634395) | 7167237 (245825) | 3649194 (122259) | 1830525 (60378) |
| Unique reflections | 15196 (2987) | 150978 (4965) | 132501 (4409) | 132666 (4382) | 132754 (4328) |
| Multiplicity | 6.9 (7.0) | 122.6 (127.8) | 54.1 (55.8) | 27.5 (27.9) | 13.8 (14.0) |
| Completeness (%) | 93.77 (71.68) | 99.97 (99.82) | 99.98 (99.86) | 99.97 (99.77) | 99.90 (99.64) |
| Mean I/sigma (I) | 10.30 (1.37) | 35.49 (1.10) | 29.63 (0.98) | 15.92 (0.81) | 13.99 (0.66) |
| Wilson B-factor | 61.99 | 65.78 | 67.58 | 59.25 | 59.02 |
| $R_{merge}$ | 0.08041 (1.147) | 0.1105 (5.697) | 0.1021 (4.39) | 0.1418 (3.556) | 0.1179 (3.148) |
| $R_{meas}$ | 0.0873 (1.238) | 0.1109 (5.719) | 0.1031 (4.43) | 0.1444 (3.621) | 0.1225 (3.267) |
| $R_{pim}$ | 0.03358 (0.4627) | 0.01004 (0.5023) | 0.01401 (0.5898) | 0.0274 (0.6818) | 0.0327 (0.8675) |
| $CC^{1/2}$ | 0.998 (0.771) | 1 (0.601) | 1 (0.508) | 1 (0.447) | 0.999 (0.345) |
| CC* | 1 (0.933) | 1 (0.866) | 1 (0.821) | 1 (0.786) | 1 (0.716) |
| *Refinement* | | | | | |
| Reflections in refinement | 14364 (2172) | 150956 (4970) | 132480 (4407) | 132652 (4382) | 132370 (4385) |
| Reflections for $R_{free}$ | 711 (101) | 7548 (249) | 6626 (221) | 6634 (219) | 6619 (219) |
| $R_{work}$ | 0.2017 (0.3587) | 0.1911 (0.2987) | 0.1972 (0.3264) | 0.2037 (0.3171) | 0.2017 (0.3400) |
| $R_{free}$ | 0.2500 (0.4169) | 0.2186 (0.3052) | 0.2260 (0.3210) | 0.2282 (0.3178) | 0.2285 (0.3677) |
| Number of non-hydrogen atoms | 2829 | 18046 | 17839 | 17927 | 17889 |
| macromolecules | 2714 | 17396 | 17379 | 17413 | 17357 |
| ligands | 90 | 262 | 265 | 254 | 250 |
| solvent | 25 | 388 | 195 | 260 | 282 |
| Protein residues | 334 | 2196 | 2194 | 2197 | 2190 |
| RMS(bonds) | 0.002 | 0.133 | 0.001 | 0.004 | 0.001 |
| RMS(angles) | 0.62 | 2.02 | 0.44 | 0.79 | 0.44 |
| Ramachandran statistics | | | | | |
| favored (%) | 98.16 | 98.02 | 99.08 | 98.94 | 97.50 |
| allowed (%) | 1.84 | 1.94 | 0.92 | 1.06 | 2.50 |
| outliers (%) | 0.00 | 0.05 | 0.00 | 0.00 | 0.00 |
| Rotamer outliers (%) | 0.97 | 0.16 | 0.05 | 0.21 | 0.11 |
| Clashscore | 2.86 | 2.87 | 2.99 | 3.51 | 3.28 |
| Average B-factor | 90.84 | 88.10 | 89.69 | 75.83 | 76.20 |
| macromolecules | 91.65 | 88.16 | 89.72 | 75.89 | 76.25 |
| ligands | 75.75 | 109.92 | 103.99 | 88.18 | 94.40 |
| solvent | 56.85 | 75.12 | 74.00 | 64.12 | 62.60 |
| Number of TLS groups | 15 | 30 | 34 | 25 | 27 |

Statistics for the highest-resolution shell are shown in parentheses.

**Size Exclusion Chromatography coupled to Multiangle Light Scattering (SEC-MALS) for assessing ternary complex formation**

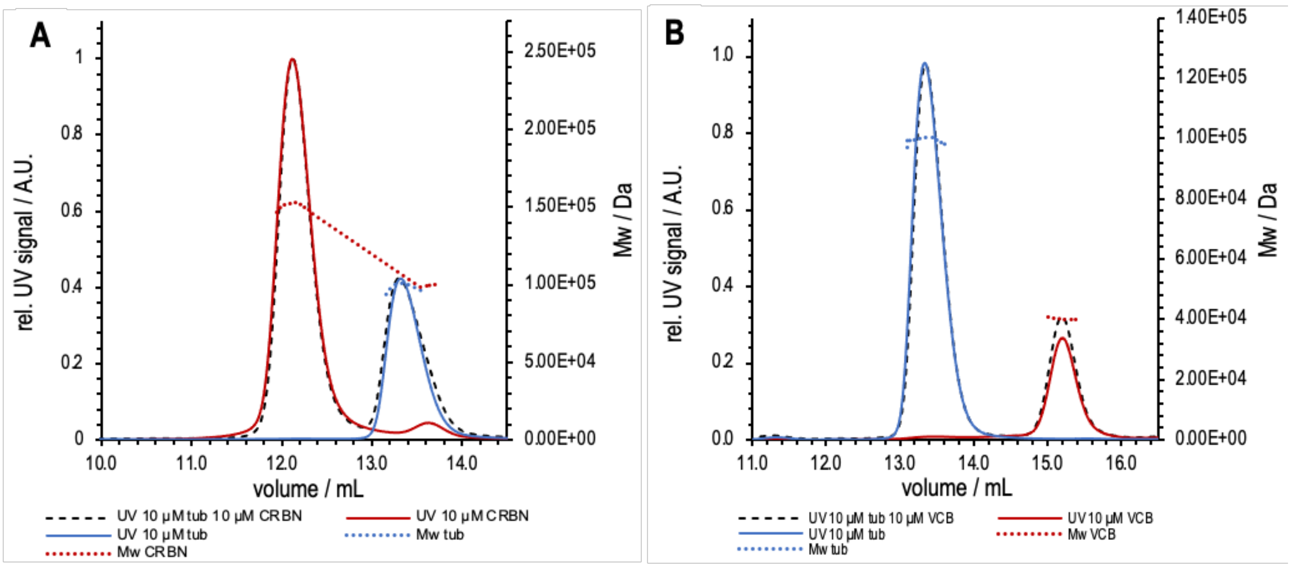

**Supplementary Fig. S5.** Reference measurements for SEC-MALS experiments. The UV signal of chromatograms obtained by injection of samples onto a Superdex 200 Increase 10/300 SEC column in BRB80 are shown for the reference measurements of 10 µM tubulin (blue), 10 µM of either CRBN or VCB (red, panels A or B, respectively) and the corresponding equimolar mixture (black dashed lines). The measured molecular weights are shown (Y2-axis on the right) as dotted lines that are colour-coded by sample.

#### Cytotoxicity assay

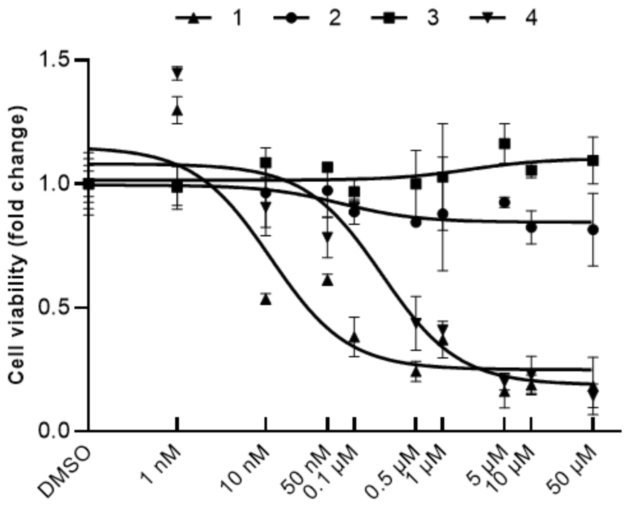

**Supplementary Fig. S6.** Dose-response curves to determine the $IC_{50}$ on cell viability for each compound, in CaCo-2 cell line.

### $^{1}H$ and $^{13}C$ NMR spectra

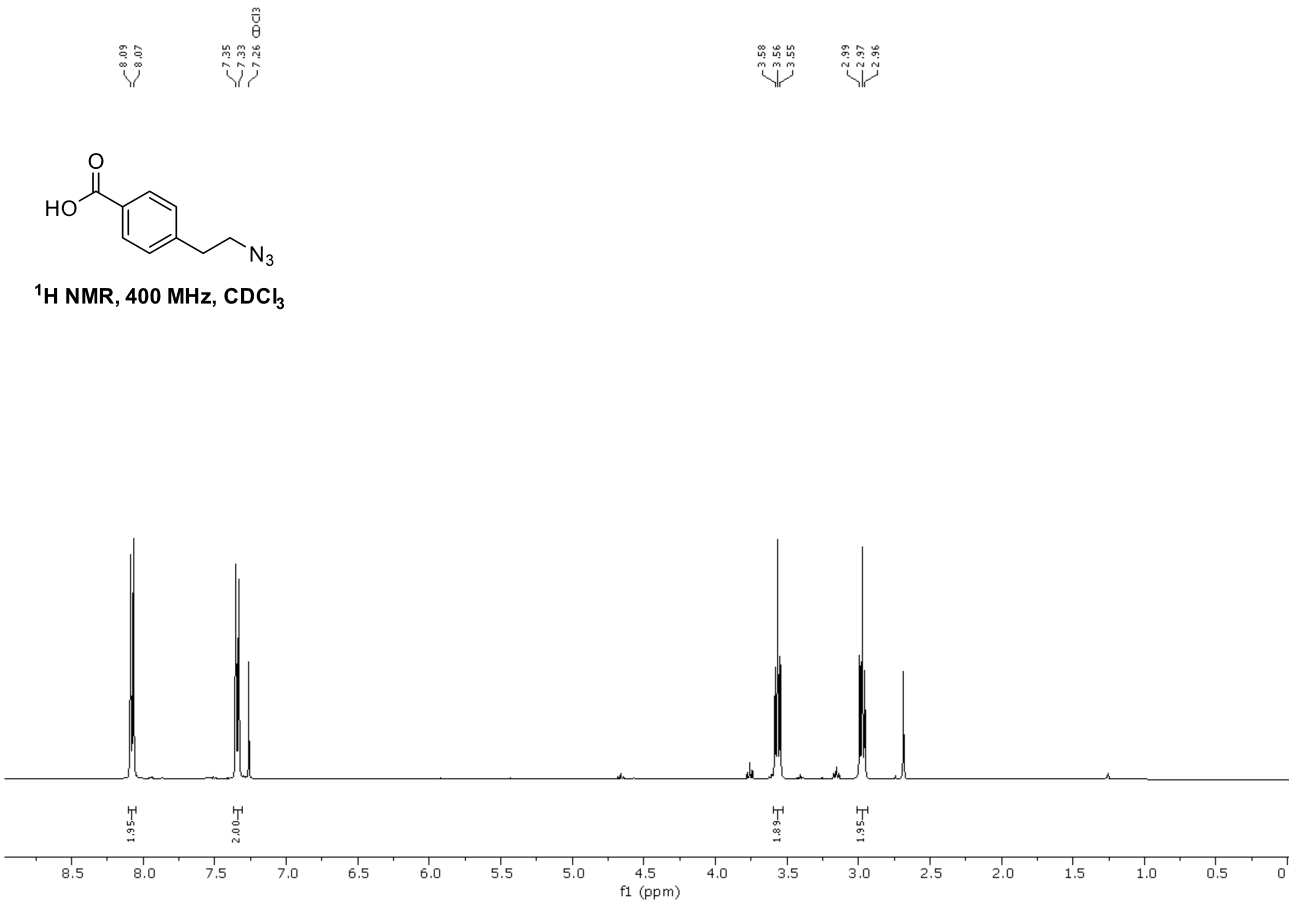

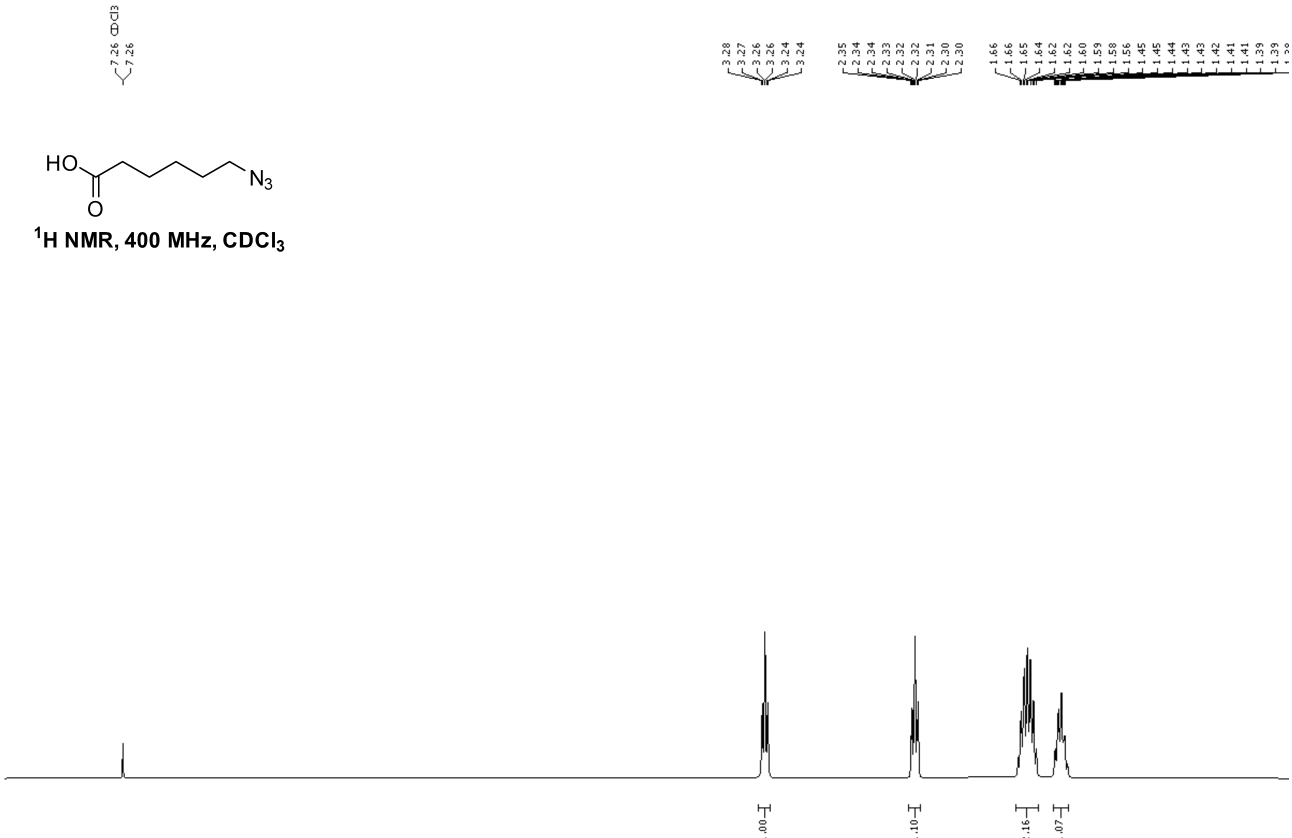

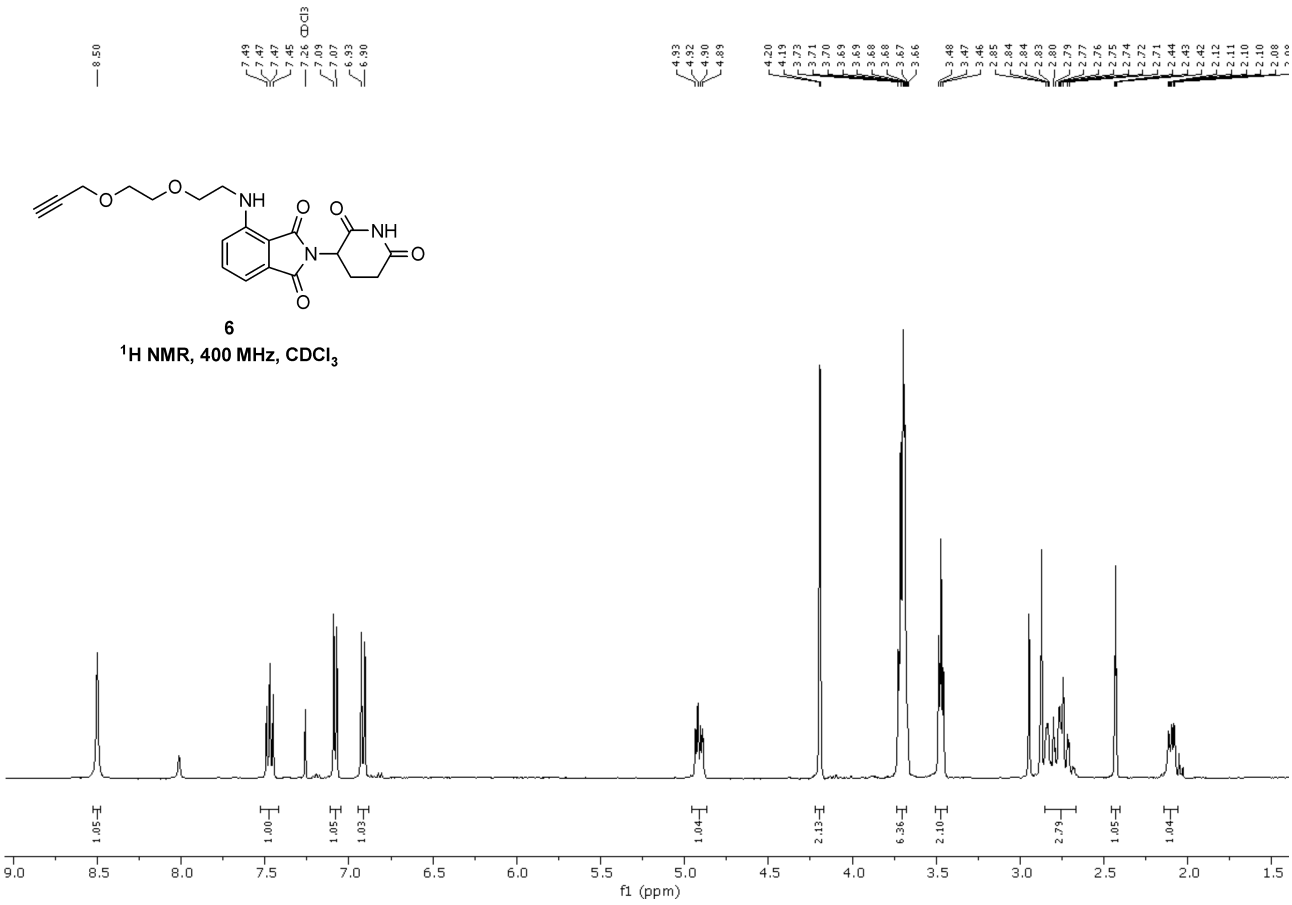
8.50
7.49 7.47 7.47 7.45 7.26 CDCl3 7.09 7.07 6.93 6.90
4.93 4.92 4.90 4.89
4.20 4.19 3.73 3.71 3.70 3.69 3.69 3.68 3.68 3.67 3.66
3.48 3.47 3.46 2.85 2.84 2.84 2.83 2.80 2.79 2.77 2.76 2.75 2.74 2.72 2.71 2.44 2.43 2.42 2.12 2.11 2.10 2.08
6
1H NMR, 400 MHz, CDCl3
1.05 1.00 1.05 1.03 1.04 2.13 6.36 2.10 2.79 1.05 1.04
f1 (ppm)

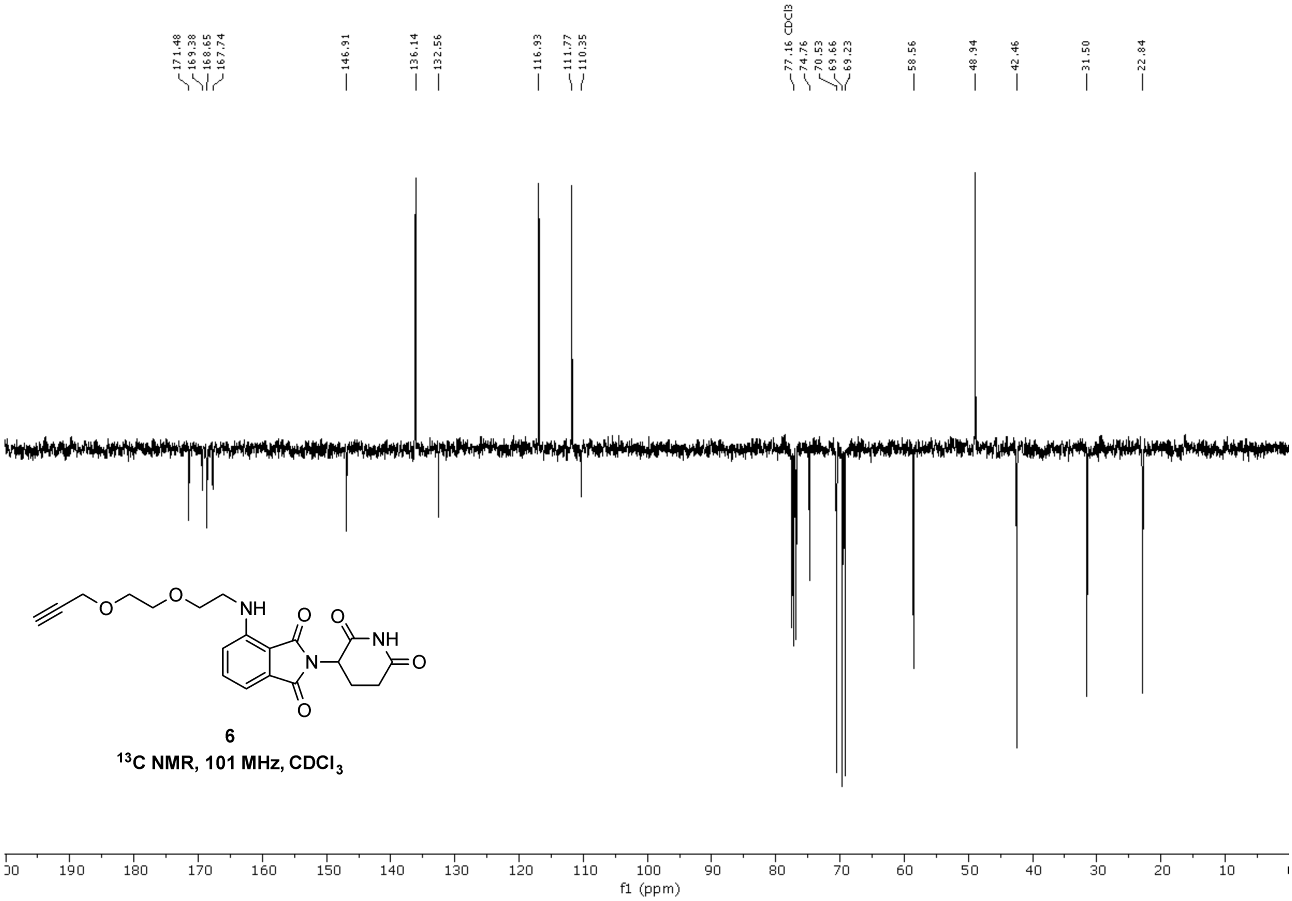
171.48 169.38 168.65 167.74
146.91
136.14 132.56
116.93 111.77 110.35
77.16 CDCl3 74.76 70.53 69.66 69.23
58.56
48.94
42.46
31.50
22.84
6
13C NMR, 101 MHz, CDCl3
f1 (ppm)

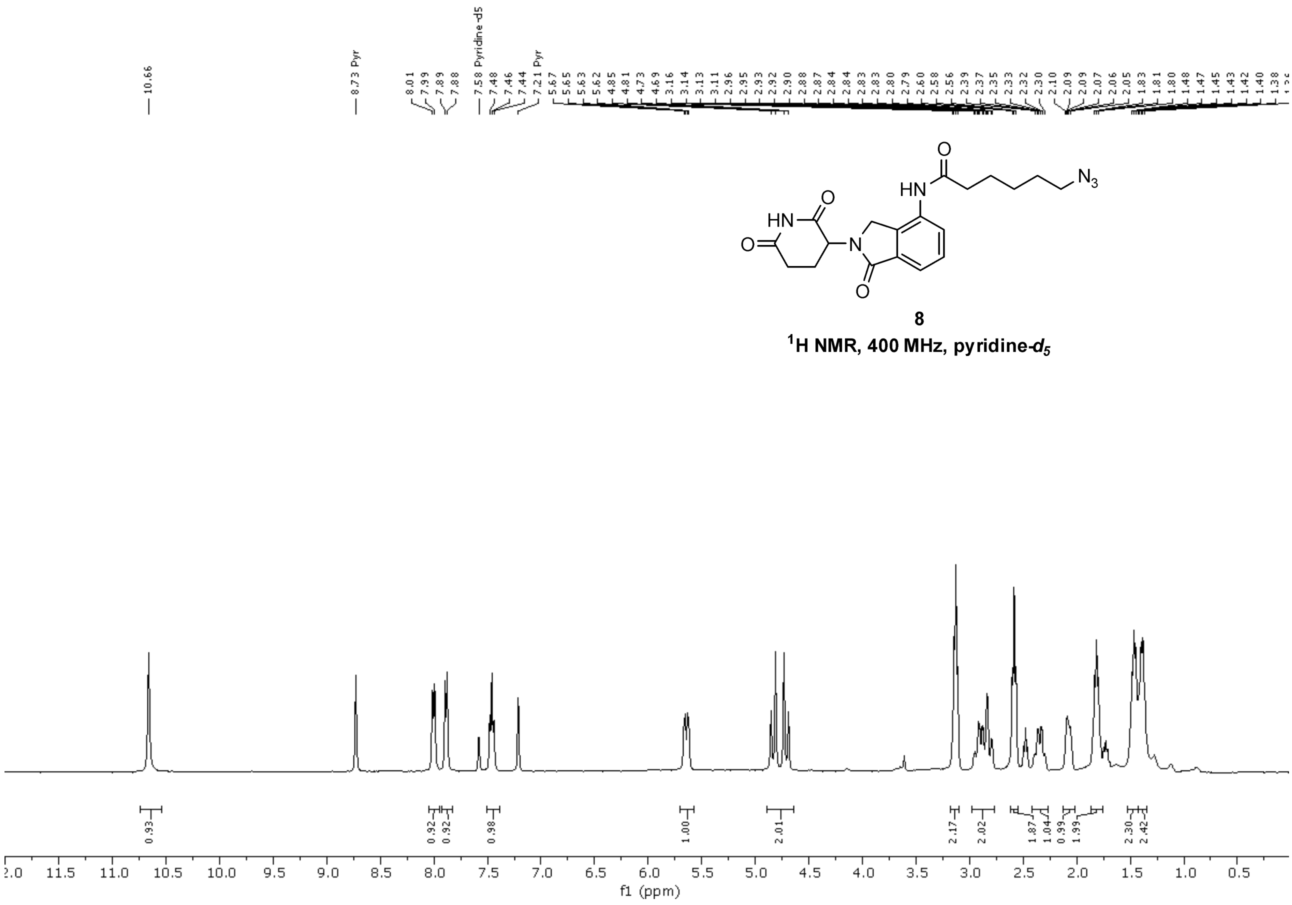

**8**

**$^{1}$H NMR, 400 MHz, pyridine-*$d_5$***

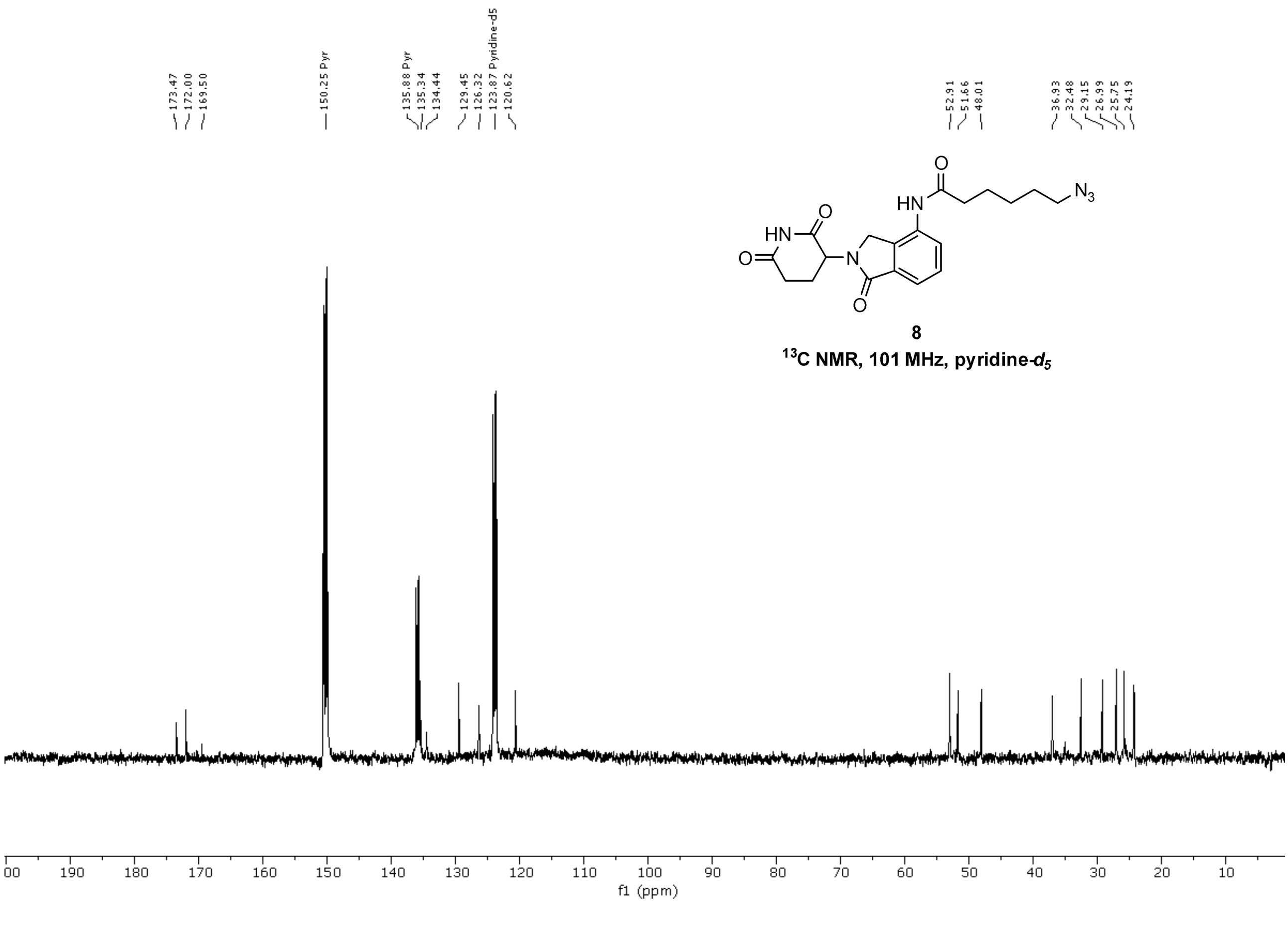

**8**

**$^{13}$C NMR, 101 MHz, pyridine-*$d_5$***

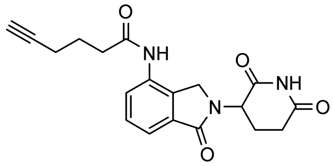

**9**

**$^{1}$H NMR, 400 MHz, pyridine-$d_5$**

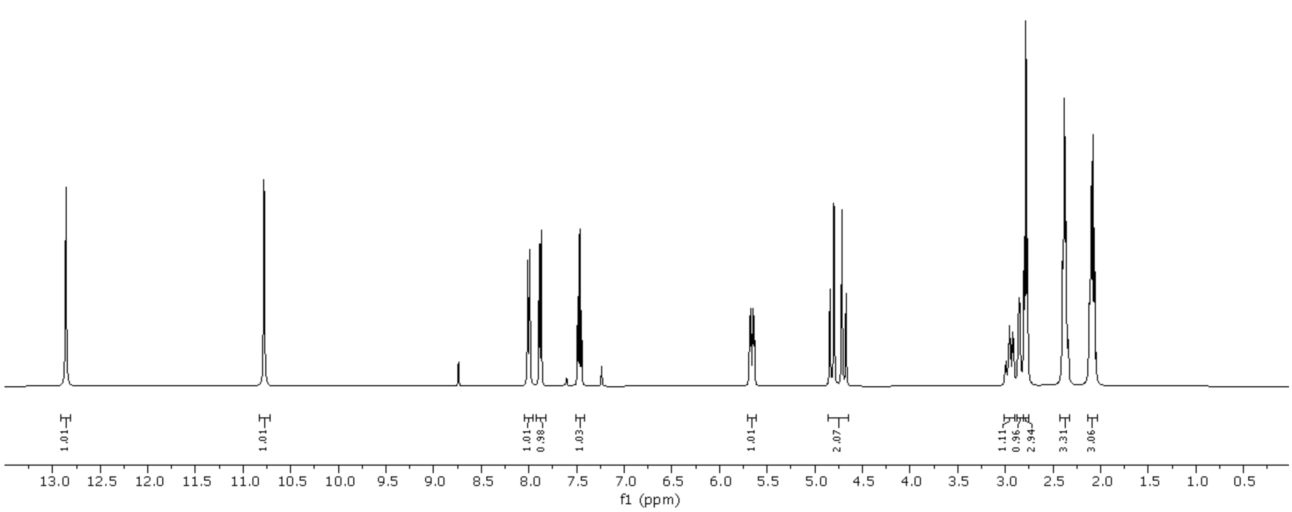

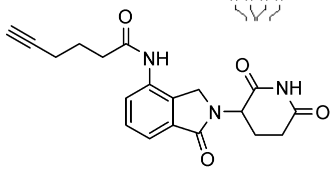

**9**

**$^{13}$C NMR, 101 MHz, pyridine-$d_5$**

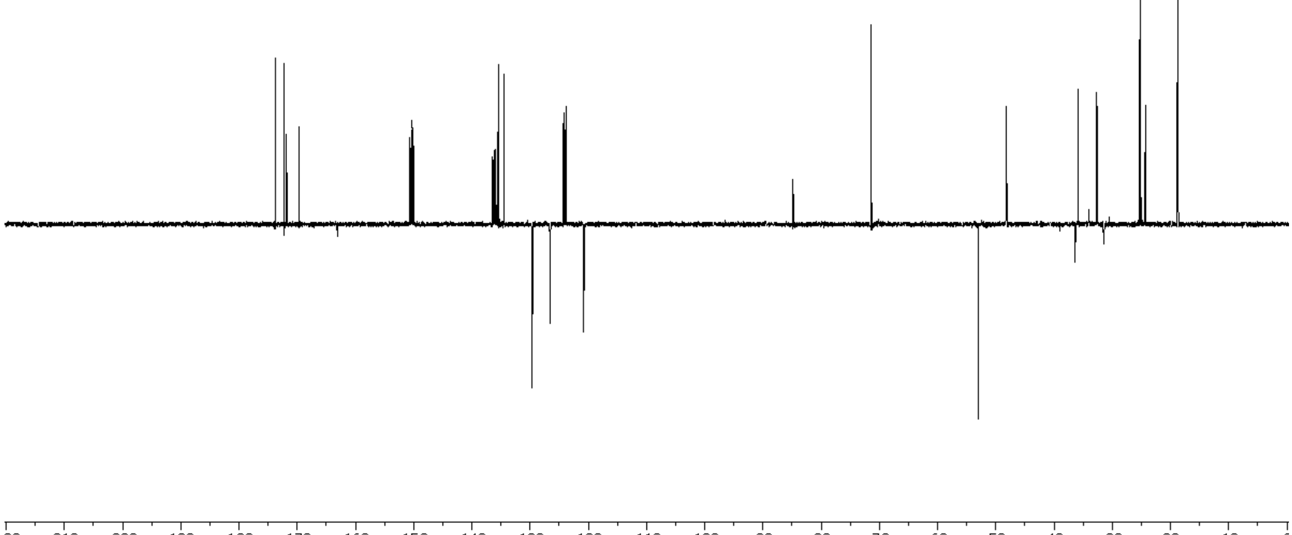

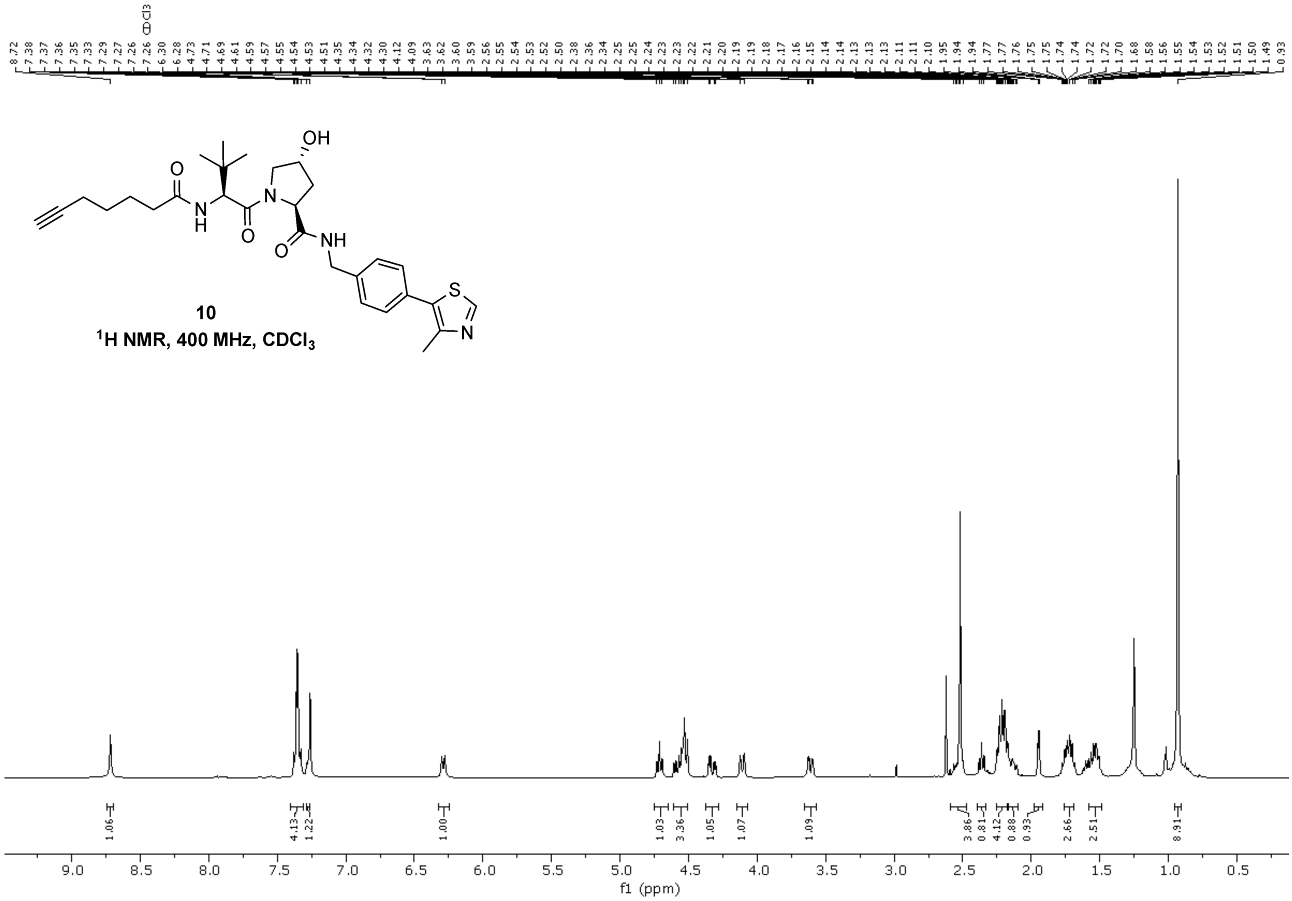

**10**
$^1$H NMR, 400 MHz, CDCl$_3$

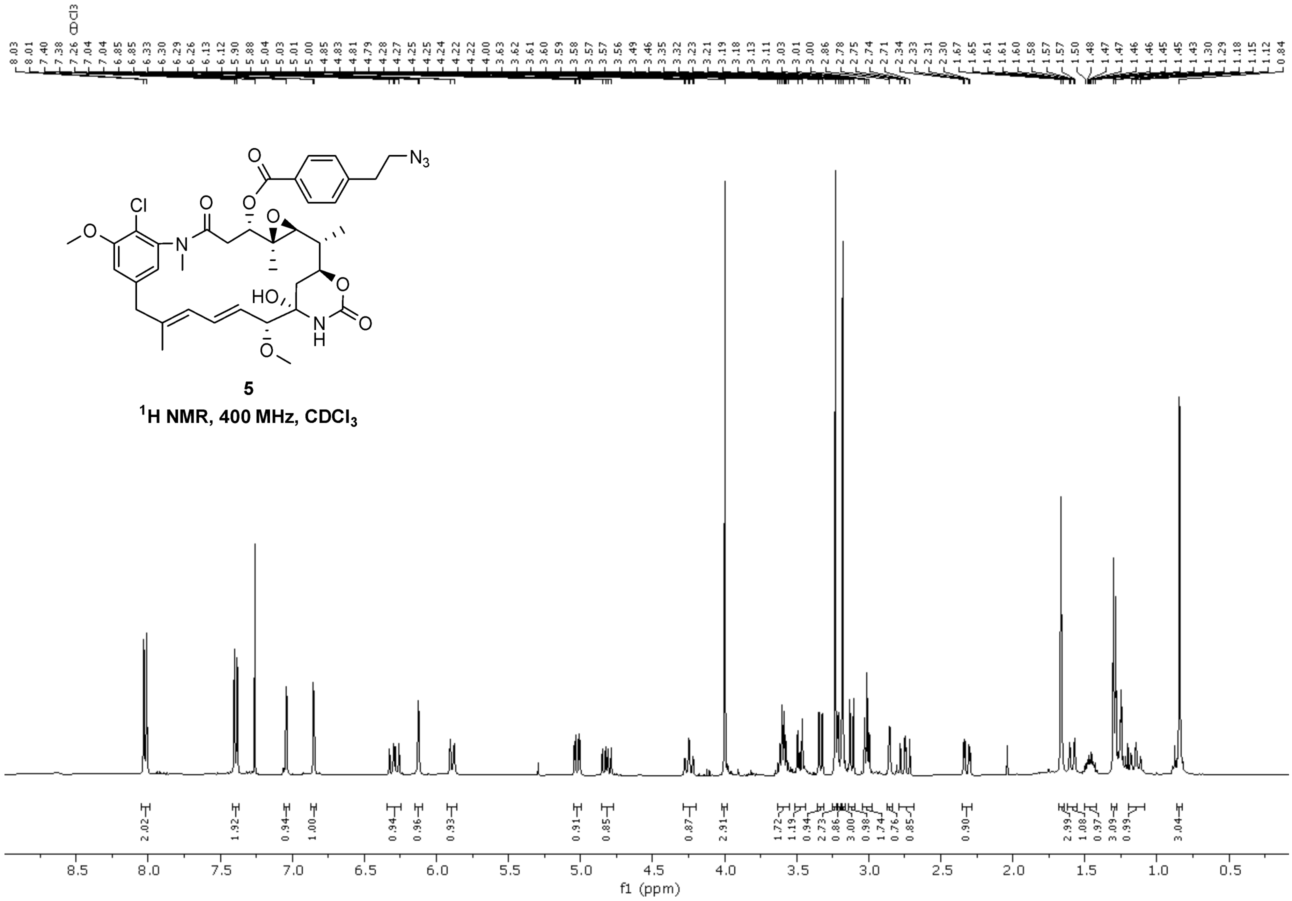

**5**
$^1$H NMR, 400 MHz, CDCl$_3$

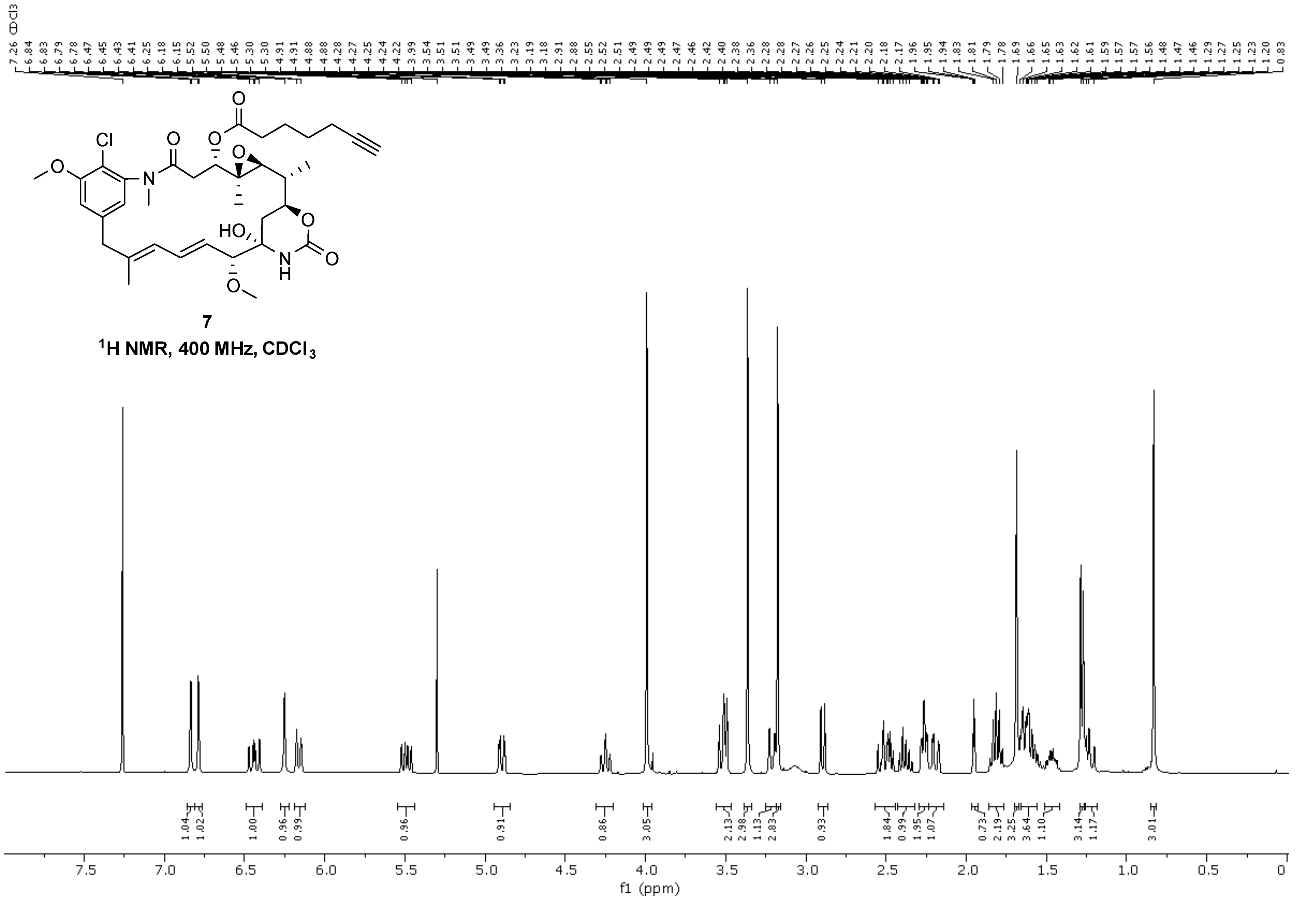
7
¹H NMR, 400 MHz, CDCl3
f1 (ppm)

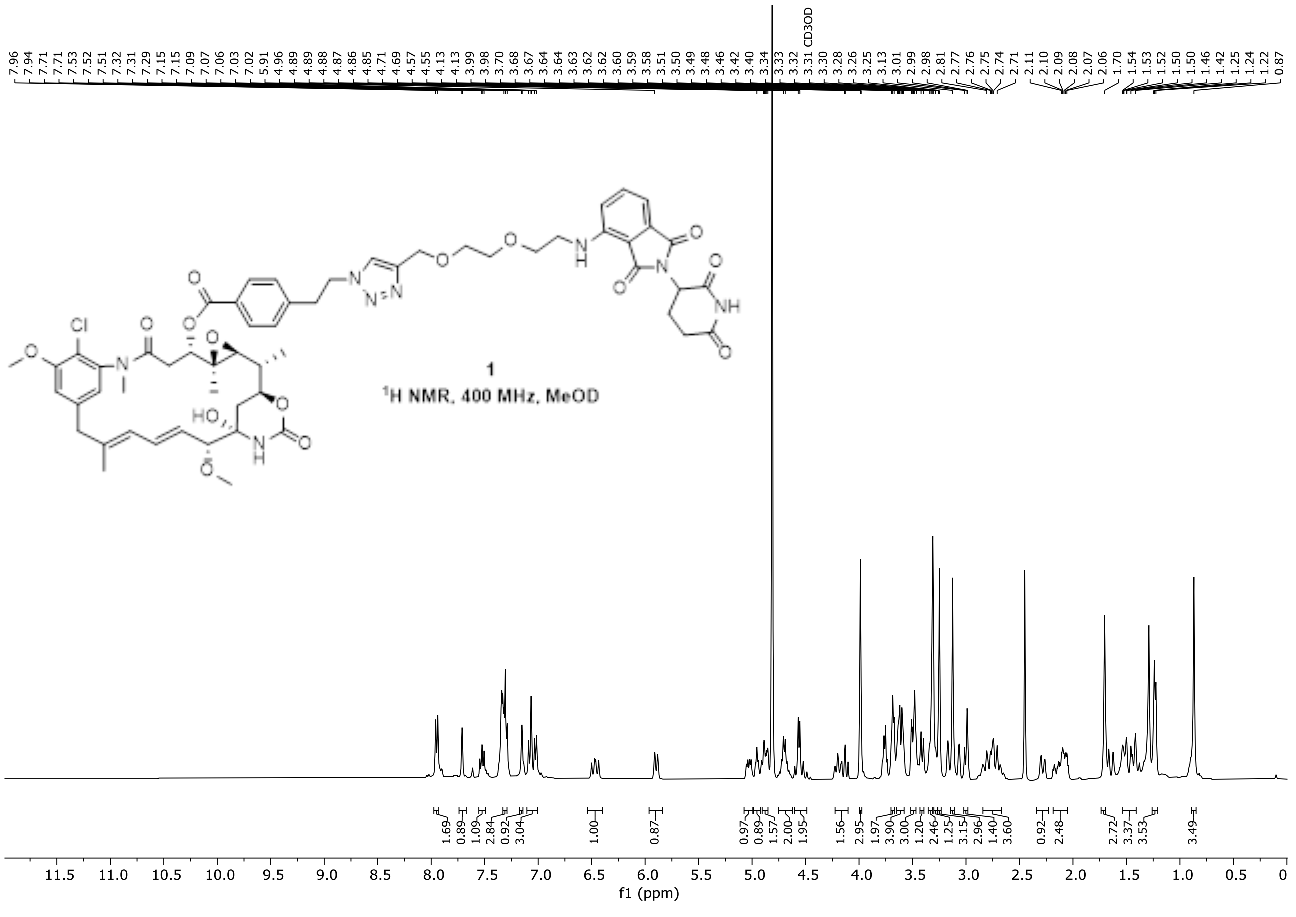
1
¹H NMR, 400 MHz, MeOD
CD3OD
f1 (ppm)

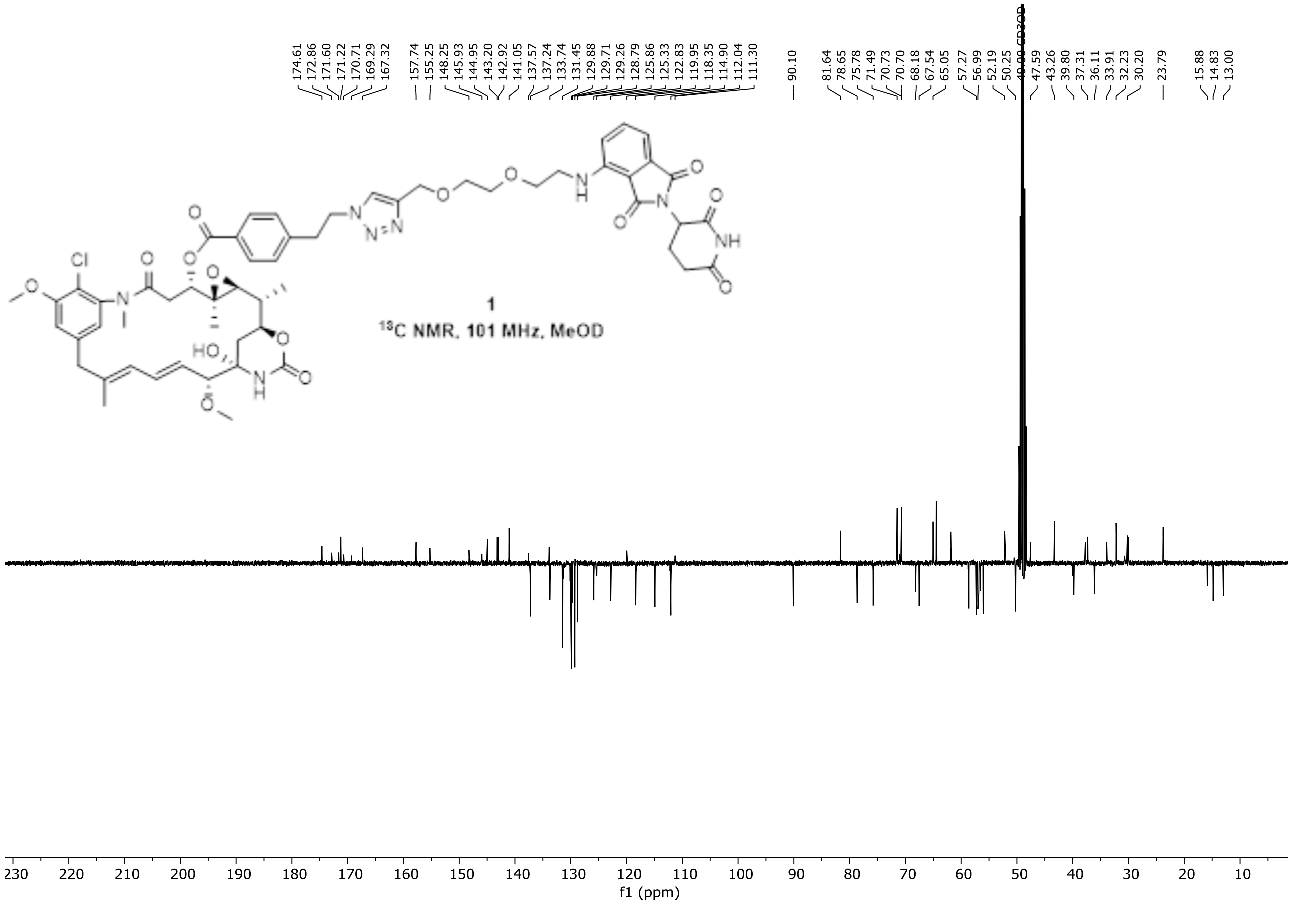
174.61
172.86
171.60
171.22
170.71
169.29
167.32
157.74
155.25
148.25
145.93
144.95
143.20
142.92
141.05
137.57
137.24
133.74
131.45
129.88
129.71
129.26
128.79
125.86
125.33
122.83
119.95
118.35
114.90
112.04
111.30
90.10
81.64
78.65
75.78
71.49
70.73
70.70
68.18
67.54
65.05
57.27
56.99
52.19
50.25
CD3OD
47.59
43.26
39.80
37.31
36.11
33.91
32.23
30.20
23.79
15.88
14.83
13.00
Cl
1
$^{13}$C NMR, 101 MHz, MeOD
f1 (ppm)

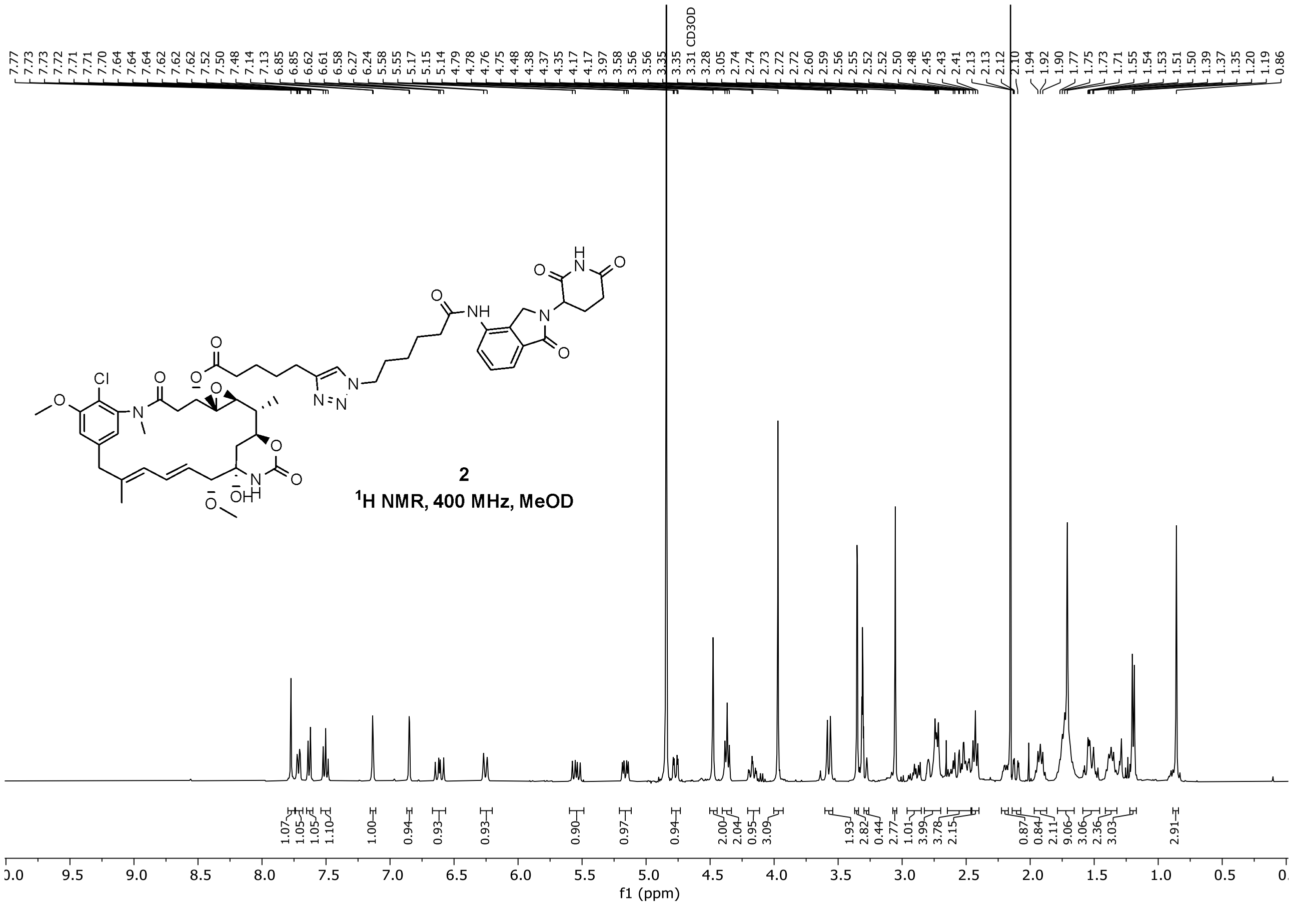
CD3OD
Cl
2
$^1$H NMR, 400 MHz, MeOD
f1 (ppm)

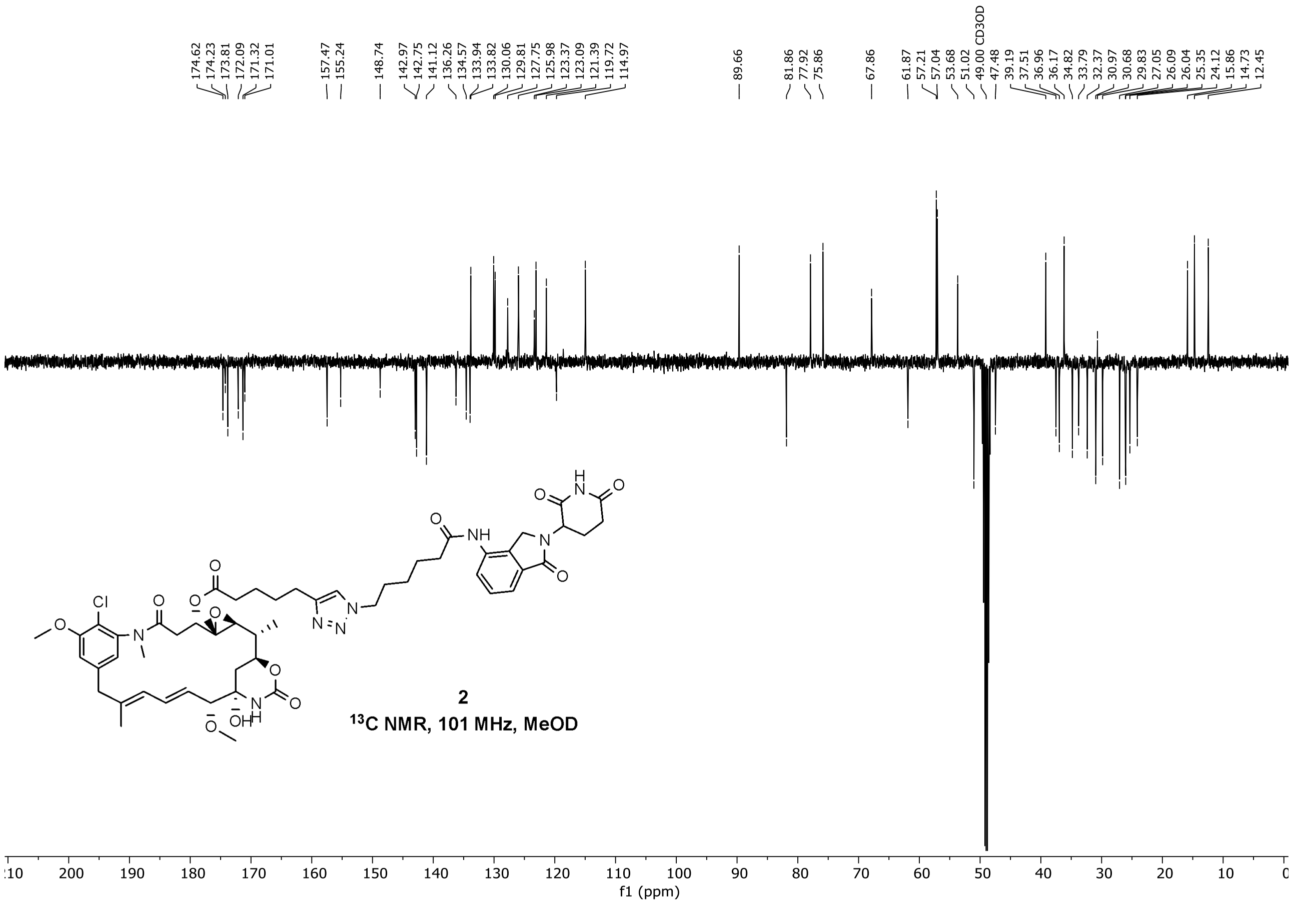
174.62
174.23
173.81
172.09
171.32
171.01
157.47
155.24
148.74
142.97
142.75
141.12
136.26
134.57
133.94
133.82
130.06
129.81
127.75
125.98
123.37
123.09
121.39
119.72
114.97
89.66
81.86
77.92
75.86
67.86
61.87
57.21
57.04
53.68
51.02
49.00 CD3OD
47.48
39.19
37.51
36.96
36.17
34.82
33.79
32.37
30.97
30.68
29.83
27.05
26.09
26.04
25.35
24.12
15.86
14.73
12.45
Cl
NH
2
$^{13}$C NMR, 101 MHz, MeOD
f1 (ppm)

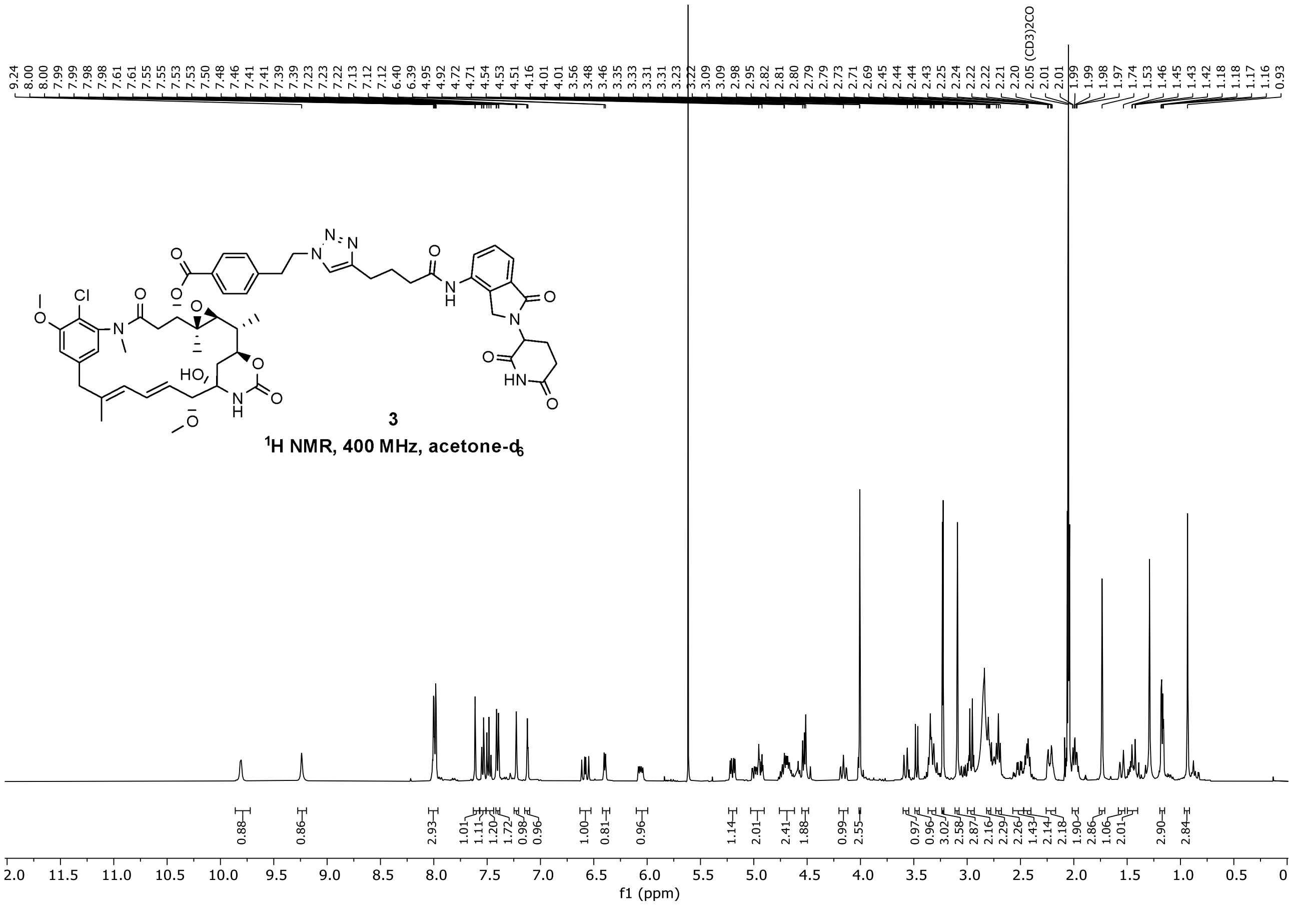
9.24
8.00
8.00
7.99
7.99
7.98
7.98
7.61
7.55
7.55
7.53
7.53
7.50
7.48
7.46
7.41
7.41
7.39
7.39
7.23
7.23
7.22
7.13
7.12
7.12
6.40
6.39
4.95
4.92
4.72
4.71
4.54
4.53
4.51
4.16
4.01
4.01
3.56
3.48
3.46
3.35
3.33
3.31
3.31
3.23
3.22
3.09
3.09
2.98
2.95
2.82
2.81
2.80
2.79
2.79
2.73
2.71
2.69
2.45
2.44
2.44
2.43
2.25
2.24
2.22
2.22
2.21
2.20
2.05 (CD3)2CO
2.01
2.01
1.99
1.99
1.98
1.97
1.74
1.53
1.46
1.45
1.43
1.42
1.18
1.18
1.17
1.16
0.93
Cl
HO
3
$^{1}$H NMR, 400 MHz, acetone-d$_6$
0.88
0.86
2.93
1.01
1.11
1.20
1.72
0.98
0.96
1.00
0.81
0.96
1.14
2.01
2.41
1.88
0.99
2.55
0.97
0.96
3.02
2.58
2.87
2.16
2.29
2.26
1.43
2.14
2.18
1.90
2.86
1.06
2.01
2.90
2.84
f1 (ppm)

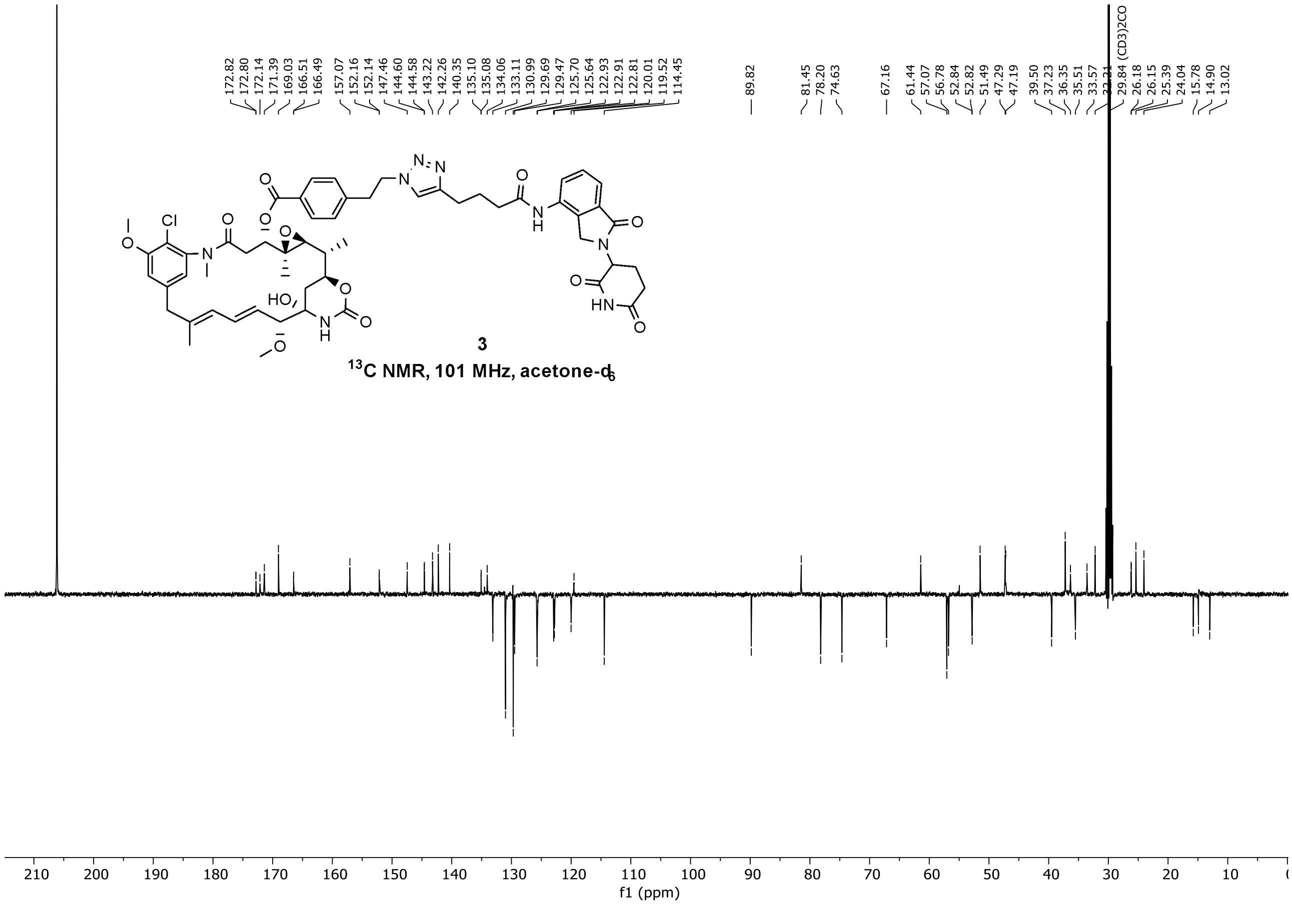
3
13C NMR, 101 MHz, acetone-d6
f1 (ppm)

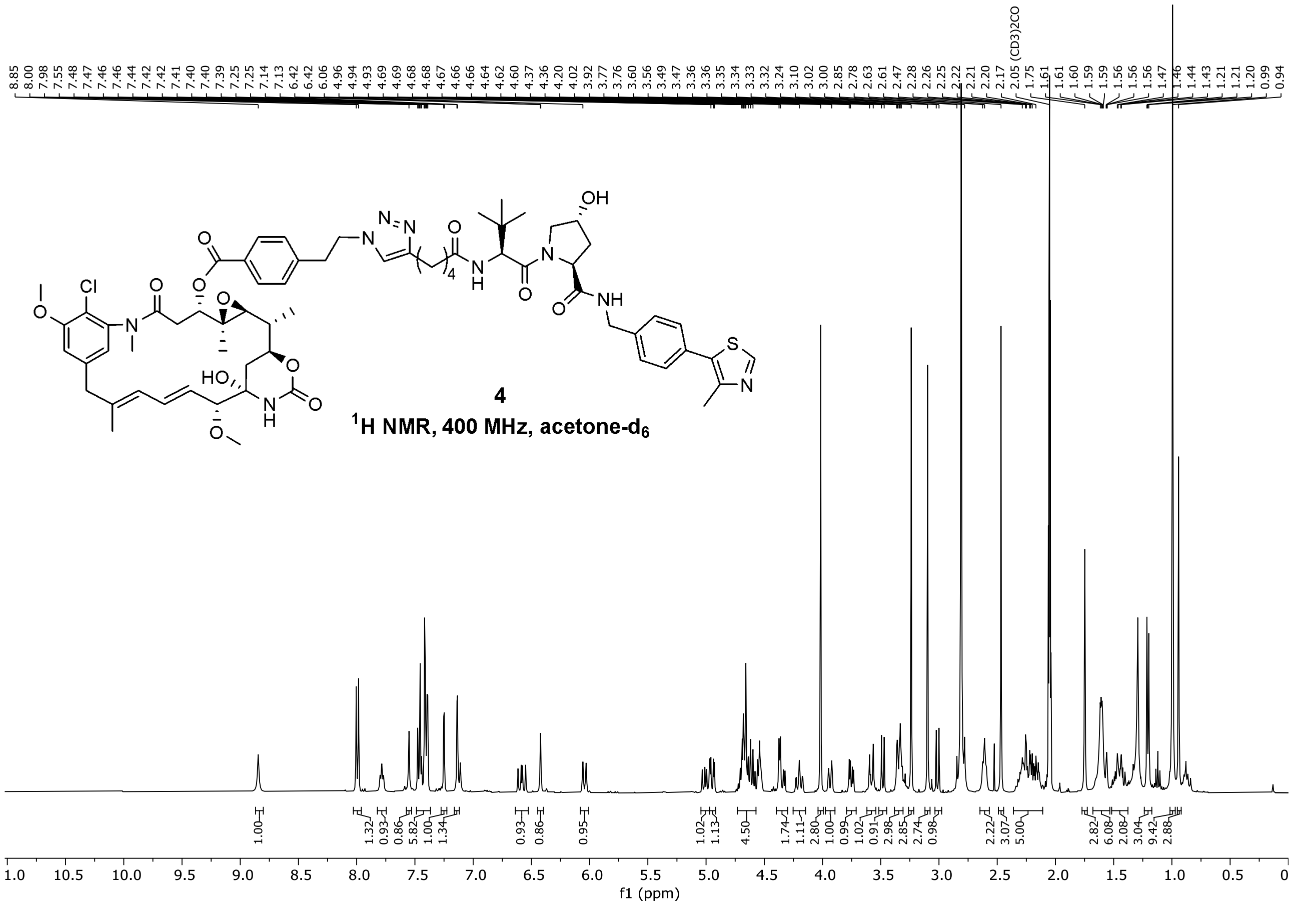
4
1H NMR, 400 MHz, acetone-d6
f1 (ppm)

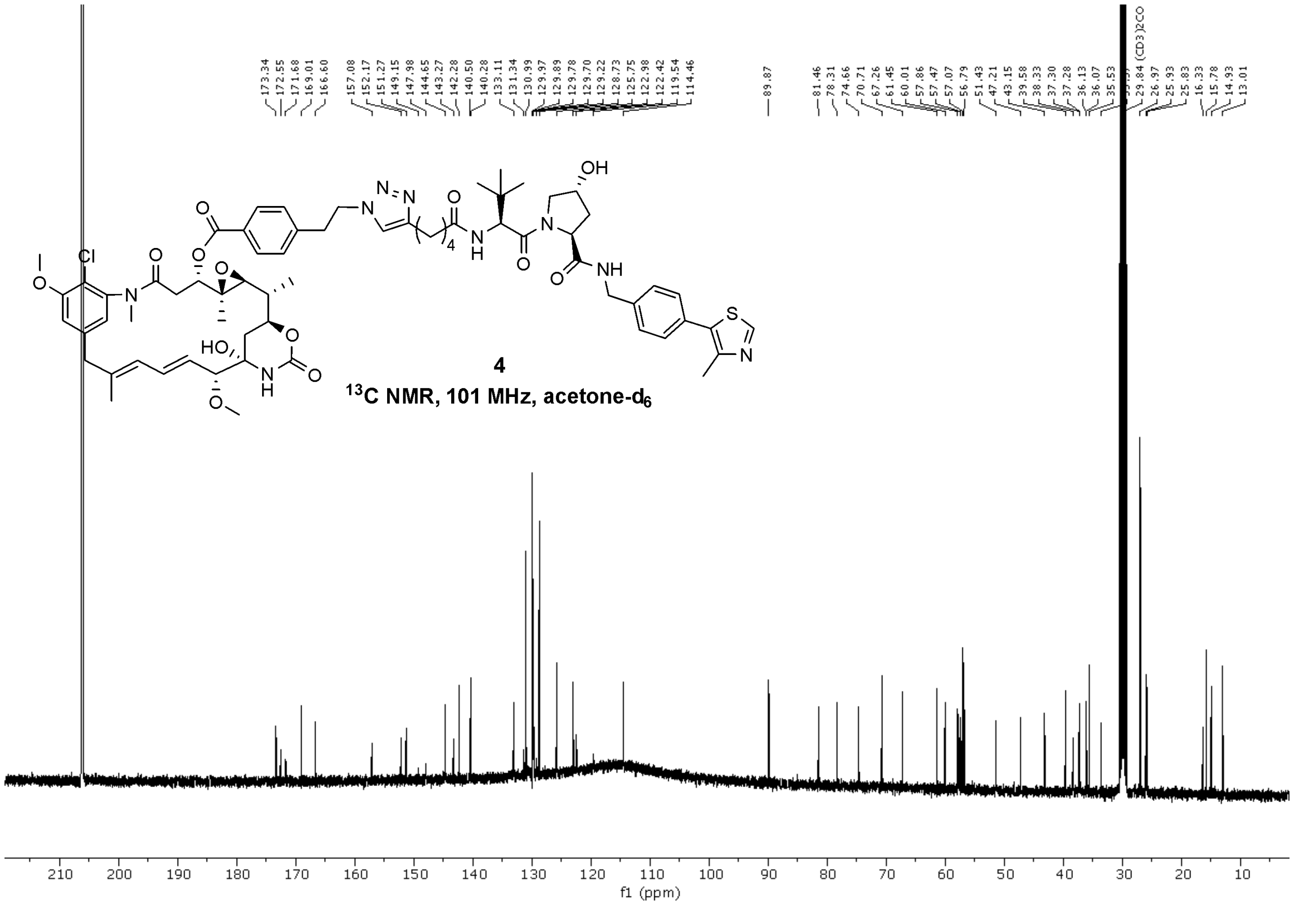
173.34
172.55
171.68
169.01
166.60
157.08
152.17
151.27
149.15
147.98
144.65
143.27
142.28
140.50
140.28
133.11
131.34
130.99
129.97
129.89
129.78
129.70
129.22
128.73
125.75
122.98
122.42
119.54
114.46
89.87
81.46
78.31
74.66
70.71
67.26
61.45
60.01
57.86
57.47
57.07
56.79
51.43
47.21
43.15
39.58
38.33
37.30
37.28
36.13
36.07
35.53
29.84 (CD3)2CO
26.97
25.93
25.83
16.33
15.78
14.93
13.01
4
13C NMR, 101 MHz, acetone-d6
210
200
190
180
170
160
150
140
130
120
110
100
90
80
70
60
50
40
30
20
10
f1 (ppm)
